# Eco-Evolutionary Optimal Carbon Allocation in a Mechanistic Crop Growth Model: Theory and Application to Wheat

**DOI:** 10.64898/2026.09.20.753045

**Authors:** Alexander J. Norton, Justin O. Borevitz

## Abstract

Carbon allocation governs how plants partition assimilates among leaves, roots, stems, and reproductive organs, directly shaping growth and yield. Most crop models represent this partitioning using fixed or empirically derived coefficients, which limits their ability to predict how allocation responds to genotype, environment, or their interaction. Eco-evolutionary optimality theory offers an alternative where the allocation strategy is allowed to emerge from the marginal costs and benefits of investing carbon in each organ. While this approach has been applied successfully to trees, it has not previously been developed for herbaceous plants. Here, we present DAESIM2-Plant, a mechanistic plant growth model that couples physiological processes including photosynthesis, stomatal conductance, plant hydraulics, and canopy radiative transfer, with an eco-evolutionary optimal allocation scheme for leaves and roots. This is applied to wheat by coupling it to a source–sink grain production module. Using a series of idealized sensitivity experiments, we show that the model reproduces well-documented patterns of plasticity in carbon allocation: diminishing returns on leaf investment as canopy closure and water limitation are approached, a shift in allocation toward roots under drier conditions, and a root:shoot response that depends jointly on soil moisture and the plant’s existing root:leaf balance. Simulating a full growing season across a gradient of soil moisture levels reveals a threshold-like response in canopy development, biomass accumulation, and grain yield, with yield constrained by the same assimilate supply that governs vegetative growth during the critical period and grain filling. This provides a promising basis for generalizing model behaviour across a wide range of environmental conditions. Evaluating these results against experimental and field trial data remains an important next step.

## 2 Introduction

Agricultural ecosystems are increasingly challenged by climate change, extreme weather, and land degradation. Simultaneously, global food production must expand to meet the demands of a growing population while minimizing environmental impact. Addressing these dual pressures requires innovative tools that enhance both productivity and resilience. Research into genetic improvements (e.g. Cooper et al., 2014) and management practices has been highly fruitful to address these pressures. Crop simulation models are central to both of these research efforts, as they help scientists understand important relationships between the management, environment and biology (Asseng et al., 2019; Hammer et al., 2014) as well as to provide predictions to support decision making.

Crop growth modeling can be traced back to efforts in the 1960s when biological and physical principles were used to describe photosynthesis mathematically (de Wit, 1965). Over time, these models have evolved and expanded to incorporate plant physiology, soil physics, hydrology, climatology, and agronomy, depending on their intended application. Today, they are widely used for crop yield forecasting, climate impact assessments (e.g. Asseng et al., 2019), and exploring genotype x environment x management (GxExM) interactions.

This diversity of applications has led to a range of modeling approaches, including: (i) Empirical models, which rely on observed data and statistical relationships; (ii) Mechanistic models, which simulate processes from first principles, often at organ or cellular scales; and (iii) Data-driven models, which use algorithms such as machine learning to identify patterns and make predictions. Hybrid approaches are also being developed (e.g. Droutsas et al., 2022; Sun et al., 2022). Empirical models are particularly useful for describing observed relationships when sufficient data are available. However, they are inherently context-dependent, often tailored to specific locations, environmental conditions, or crop types, and can be unreliable when extrapolated to new conditions. Data-driven approaches can achieve high predictive accuracy (Leng and Hall, 2020), but they typically lack explanatory power regarding underlying mechanisms. Consequently, they are poorly suited for hypothesis-driven exploration, such as investigating causal processes or conducting “what-if” analyses, which limits their interpretability and utility in exploring GxExM interactions. In contrast, mechanistic models offer a theoretical framework in which outcomes (e.g., crop yield) are explicitly linked to physical and biological principles. This allows outcomes to be traced back to underlying processes, enabling deeper investigation of G×E×M interactions (e.g. Wu et al., 2019) and the identification of emergent relationships .

Mechanistic models aim to simulate biological processes using equations grounded in physical and biological principles. These formulations typically include parameters that have biophysical meaning and explicit units. However, in practice, many components within mechanistic models are still represented using empirical parameters or even empirical functions derived from limited observational data. This may be due to limited theoretical understanding, limited data availability, or both. This blending of approaches reduces a model’s capacity to explore causal mechanisms or evaluate complex GxExM interactions. For instance, a model may represent photosynthesis mechanistically but the partitioning of these assimilates to different organs may be fixed or determined using empirical functions. This is the case for many widely used crop models including WOFOST (de Wit et al., 2019) and APSIM (Yang et al., 2023). Thus, changes in photosynthesis either by environmental conditions or assigning different traits (e.g. photosynthetic capacity) will have no effect on the plants strategy for growth. Using fixed or empirical coefficients embeds assumptions about plant behavior that are not explicitly testable nor supported by data. As Connor and Fereres (1999) noted, such models often “apply a good part of the answer” rather than propose hypotheses that can be evaluated and refined.

Carbon allocation is a critical component of crop models, as it governs how assimilates are distributed among organs and functions, directly influencing relative growth rates and yield. It is not a single process but the outcome of several distinct ones including the transport, storage and utilisation of assimilates, governed jointly by the supply of assimilates from photosynthesis and the capacity of different organs to use that supply (Cannell and Dewar, 1994; Brüggemann et al., 2011). Although significant progress has been made in our understanding of the molecular mechanisms that regulate this partitioning, including genes, transport pathways and signalling (Fernie et al., 2020; Rosado-Souza et al., 2023; Sonnewald and Fernie, 2018), this understanding is not yet sufficient to develop a carbon allocation model that is both parsimonious and generalisable. Nevertheless, carbon allocation is involved in multiple feedback processes that shape plant behaviour over its life cycle. Without a robust representation of these dynamics, even well-developed representations of other processes may fail to produce reliable outcomes for many crop modeling applications.

There is strong empirical evidence that plants dynamically adjust carbon allocation in response to resource limitations, demonstrating plasticity in carbon allocation (e.g. Shipley and Meziane, 2002). For example, under water stress, plants reduce shoot growth and leaf area while enhancing water-use efficiency and reallocating carbon belowground (Vandoorne et al., 2012; Chandregowda et al., 2023). Rooting depth has been shown to vary with local soil hydrology, reflecting adaptability to climate, waterlogging, and moisture profiles (Fan et al., 2017). These shifts support recovery and resource acquisition under adverse conditions. The root:shoot ratio, a commonly used indicator of biomass partitioning, also varies with soil texture and nutrient availability, with clay-rich soils typically reducing investment in root biomass due to hydraulic considerations (Poeplau and Kätterer, 2017; Qi et al., 2019; Ding et al., 2025). Such plastic responses are qualitatively consistent with an optimal allocation hypothesis (McCarthy and Enquist, 2007), which posits that plants adjust the partitioning of assimilates in response to resource limitations (Bloom et al., 1985; Chapin et al., 1987).

However, resource limitation is not purely an environmental property. It is jointly determined by the environment and the traits of the plant itself. What is considered “limiting” for a given plant depends on its genetically determined phenotype, so two genotypes in the same environment can experience different degrees or even different kinds of resource limitation. Recent work has shown that a wheat genotype with a greater capacity for root growth under drought maintains higher root water influx and transpiration than a genotype with a more conservative allocation strategy under the same water deficit (Bacher et al., 2022), supporting greater growth and yield (Bacher et al., 2022). Similarly, genotypic variation in the fraction of biomass allocated to leaves in sunflower is directly linked to whole-plant transpiration efficiency, independent of leaf-level physiological traits (Velázquez et al., 2017), whereby larger leaf area increases evaporative demand for a given water supply. Carbon allocation therefore reflects an interplay between trait and environment.

Models that rely on fixed allocation coefficients or empirical functions risk overlooking both the environmental drivers of allocation and their dependence on plant traits. Some existing modeling approaches do capture elements of these resource limitations. Functional balance theory (also called functional equilibrium), for instance, predicts that shoot growth responds to the light or CO_2_ available per unit of root, while root growth responds to the water or nutrients available per unit of shoot, such that allocation shifts to balance the two (Poorter and Nagel, 2000). This qualitatively predicts the direction of allocation shifts under light, water, or nutrient limitation, but it does not provide a mechanistic, trait-resolved basis for predicting the allocation coefficients themselves, nor does it naturally accommodate multiple, simultaneous resource limitations. Other modeling approaches capture some of this behavior empirically, through allocation coefficients that vary by growth stage or that are modified as a function of nutrient or water status, but such relationships are typically calibrated for specific crops, sites, or conditions and do not generalize well beyond them. An integrated theory that captures both the environmental and trait-dependent determinants of resource limitation, while remaining broadly generalizable across species and conditions, has not been developed for crop models. Recent applications of eco-evolutionary optimality have proven highly useful in describing plant behaviour without the need for highly parameterized functions (Franklin et al., 2020; Harrison et al., 2021). For example, Franklin (2007) used an optimality theory to predict how nitrogen limitation controls the response of forest growth to elevated atmospheric carbon dioxide, and Potkay et al. (2021) developed an eco-evolutionary optimality theory for carbon allocation in trees that provided good predictions of tree biomass partitioning across tree sizes, water limitations, elevated atmospheric carbon dioxide, and response to pruning. While eco-evolutionary optimality has been applied to allocation in trees, an equivalent approach has not been developed for grasses or annual crops, which differ substantially from trees in growth form, allocation strategy, and life history.

Here, we describe a new crop growth model that integrates mechanistic representations of plant growth and development with an eco-evolutionary optimality theory for carbon allocation to leaves and roots. This applies an optimal-path formulation in which the target itself shifts continuously with conditions, rather than a fixed target allometry. The aims are: (i) to describe the model structure and the theoretical basis of its novel components, (ii) to demonstrate, through idealised sensitivity simulations, how the optimality-based allocation framework reproduces commonly observed patterns of plant growth and biomass partitioning under varying resource conditions, and (iii) to illustrate how the interplay between environmental conditions and plant traits jointly determines carbon allocation outcomes, and hence growth and yield.

## 3 Methods

The Dynamic Agro-Ecosystem Simulator (DAESIM) was developed by Taghikhah et al. (2022) and includes modules for carbon cycling (plant, litter, and soil), water cycling, nutrient cycling (nitrogen, phosphorus), soil erosion, and the effects of livestock on plants and soils. This first version of DAESIM was designed primarily for summarizing and assessing natural capital in managed landscapes. While many of the key components of managed landscapes are included, the specific processes and interactions are represented with a fairly simple stock and flow model structure, which lacks many known dependencies on environmental conditions.

Here, we describe a new process-based crop simulation model developed for the second version of DAESIM, DAESIM2. This crop model is designed to represent crop structure, function and development with core physiological and biophysical processes. The focus of this new crop model is upon plant carbon and water cycling. Wherever feasible, the model integrates carbon and water cycling using mechanistic formulations and trait-based parameters, which includes the key environmental drivers. The processes include photosynthesis, stomatal conductance, plant hydraulics, plant respiration, canopy radiative transfer, crop development, grain production and dynamic carbon allocation. Key features of the model include:

- Physiological realism: Core processes such as photosynthesis, respiration, stomatal conductance, plant hydraulics and canopy radiative transfer are represented mechanistically.
- Optimality theory for carbon allocation: Allocation to leaves and roots is guided by principles of eco-evolutionary optimality, not fixed empirical rules.
- Grain production: Grain production is described using a source-sink approach, where the sink determined by critical periods and the source determined by the flow of new assimilates and retranslocation of stored assimilates in the stem.

The model currently includes implementations for two key crop types: Wheat and canola. However, most processes are generalizable across most annual and perennial crop types and may be updated in future with appropriate parameterization and crop-type specific grain production modules.

### 3.1 Model Description

DAESIM2-Plant combines physiological sub-models that were each selected on three broad criteria: (i) generalizability; (ii) process completeness; and (iii) a track record of testing and validation across species and scales. A full model description is provided as a supplement, including the mathematical formulations for each process and the relevant references.

#### 3.1.1 Plant Structure

The model is designed to represent a population of plants with horizontally homogenous traits and structure. The model assumes a horizontally uniform canopy albeit with some important features, such as shading within the canopy and foliage clumping effects. The plant is represented by four carbon pools: leaves, roots, stems and grain/seed. The plant canopy is discretized into n layers. Each layer has user-defined optical properties, physiological parameters (e.g. Vcmax) and structure (LAI, SAI, height).

The soil is discretized into m layers. Each layer has user-defined physical (e.g. bulk density, sand/silt/clay) and hydraulic properties (e.g. saturated soil water content, air-entry value of hydrostatic water potential, saturated hydraulic conductivity).

#### 3.1.2 Plant Coupled Physiological Processes

Canopy photosynthesis follows the steady-state C3 biochemical model of Farquhar et al. (1980), with temperature-dependent kinetics represented using standard Arrhenius, peaked-Arrhenius, and Q10 formulations (Medlyn et al., 2002). This model was chosen because its parameters (e.g. maximum carboxylation rate, *V*_*cmax*_; maximum electron transport rate, *J*_*max*_) correspond directly to measurable leaf biochemical traits, allowing genotypic and environmental variation in photosynthetic capacity to propagate through the model. It has also been validated extensively from the leaf to the global scale (von Caemmerer, 2000; Yin et al., 2021).

Stomatal conductance is represented using the unified optimal stomatal conductance model of Medlyn et al. (2011, 2012), implemented following Duursma (2015). This formulation was preferred over alternative empirical stomatal models (e.g. Ball–Berry-type formulations) because it is itself derived from optimality theory whereby stomata are assumed to regulate carbon gain against water loss, providing conceptually consistent with the optimality-based approach to carbon allocation used elsewhere in the model. Its key parameter, *g*_1_, has a direct physiological interpretation as the marginal water cost of carbon gain, and the model has been shown to reconcile optimal and empirical approaches to stomatal behaviour across a wide range of species and biomes (Medlyn et al., 2011), supporting its generalizability. We additionally incorporate down-regulation of stomatal conductance under declining leaf water potential using the Tuzet model (Tuzet et al., 2003), which extends the framework to represent soil moisture limitation and drought responses explicitly.

Plant hydraulics are represented as a one-dimensional, steady-state pathway from soil to leaf, following Darcy’s law and Ohm’s law analogy for hydraulic conductances in series (Bonan et al., 2014). Leaf water potential is solved so that water supply from root uptake balances water loss through transpiration (Meinzer, 2002; Bonan et al., 2014). This approach was chosen because it explicitly links plant water use to root and xylem hydraulic traits and to prescribed soil moisture, rather than assuming a fixed or empirically-derived water stress response; this allows drought responses to emerge from the interaction of soil conditions, root distribution, and hydraulic traits, rather than being imposed. Formulations of this kind are well established in land-surface and ecosystem models (Williams et al., 2001; Duursma et al., 2008; Bonan et al., 2014) and have been applied specifically to agricultural species including wheat (Ranawana et al., 2021).

Canopy radiative transfer is represented using a multi-layer two-stream approximation (Bonan et al., 2018, 2021), which partitions incident radiation into direct and diffuse components across an arbitrary number of canopy layers with distinct structural and optical properties. This was chosen in preference to simpler big-leaf or two-big-leaf schemes because those simpler approaches are known to introduce biases in simulated carbon, water and energy fluxes (Luo et al., 2018; Wang and Frankenberg, 2022), particularly for canopies with varied vertical structure, a relevant consideration for crops, whose canopy structure changes across developmental stages (Hosoi et al., 2009; Li et al., 2015).

The vertical distribution of root biomass with soil depth follows a logistic dose-response function (Fan et al., 2016), chosen because it has been parameterized and evaluated for a wide range of crop types, including wheat and canola, and captures the exponential-to-near-linear decline in root density with depth that is consistently observed empirically (Fan et al., 2016). This vertical root distribution, combined with the soil-to-root hydraulic conductance model above, allows water uptake to reflect realistic interactions between root distribution and the vertical soil moisture profile, rather than treating the soil profile as a single undifferentiated pool.

#### 3.1.3 Plant Carbon Balance and Respiration

Carbon allocation operates on the assimilate pool remaining after respiratory costs have been met, so the plant carbon balance and respiration are first described here. The rate of change of total plant carbon is:

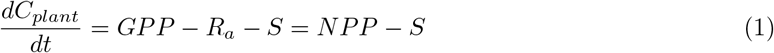

Where *R*_*a*_ is total autotrophic respiration and *S* is the litter production (senescence) rate. Autotrophic respiration follows the growth-and-maintenance paradigm (McCree, 1970; Amthor, 2000), in which all respiratory costs are attributed to either maintaining existing live biomass or constructing new biomass. Maintenance respiration, *R*_*m*_, is the sum of leaf mitochondrial respiration (scaled with photosynthetic capacity, following its shared dependence with *V*_*cmax*_ on leaf nitrogen) and root maintenance respiration, which is proportional to root biomass and temperature-sensitive following a *Q*_10_ response. Growth respiration is represented as a fixed fraction of the assimilate remaining after maintenance costs are met, GPP - Rm, reflecting the assumption that maintenance of existing tissue is prioritized over new growth. Net primary productivity is therefore:

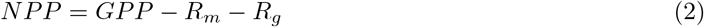

This maintenance-first structure has a direct consequence for carbon allocation such that the quantity GPP-Rm, or net carbon profit, is what the optimal allocation scheme (Section [Optimal Carbon Allocation]) evaluates when weighing the marginal benefit of investing in leaves versus roots, so respiratory costs already incurred in maintaining existing tissue are accounted for before any new allocation decision is made.

#### 3.1.4 Optimal Carbon Allocation

A central challenge in modeling plant growth is deciding how much carbon a plant should invest in each major organ (leaves, roots, stem, or grain) at any given time, and how that investment should shift as conditions change. One way of framing this problem, drawing on an economic analogy, treats the plant as an agent allocating a limited resource (carbon) among competing investments (organs), each of which returns a different, and time-varying, benefit (Bloom et al., 1985). Eco-evolutionary optimality theory extends this idea by proposing that, over evolutionary time, plants have been selected to allocate carbon in a way that maximizes some measure of long-term fitness or carbon gain, given the constraints imposed by their traits and environment (Franklin et al., 2020; Harrison et al., 2021). Rather than prescribing how allocation should respond to any particular condition, this approach allows the allocation strategy itself to emerge from the physiological processes already represented in the model.

In DAESIM2-Plant, the governing principle is that plants allocate carbon in a way that maximises the potential for net carbon profit. This defines the so-called ‘fitness proxy’. Here, this is applied to the leaves and roots, while allocation to the stem and grain are governed by alternative formulations. The carbon allocation to leaves and roots follows an instantaneous optimal trajectory principle, modified from Potkay et al. (2021). This is a so-called optimal path, or optimal trajectory, approach (Buckley and Roberts, 2006; Caldararu et al., 2020). Rather than calculating a global optimum target allocation, the model determines, at each time step, the direction in which the leaf and root carbon pools should be adjusted to increase potential net carbon profit. This target may shift continuously and need not be fully reached at any given time. We discuss the implications of this distinction, relative to more traditional optimality models, in Section 5.1.

At each time step, the fraction of new assimilate allocated to organ *k*, given by *u*_*k*_, is proportional to the ratio of the marginal carbon gain to the marginal carbon cost of investing in that organ:

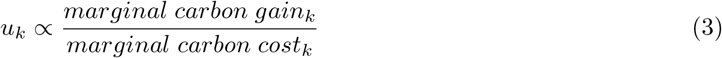

The marginal gain determines the net carbon profit that would be returned if there were one unit of investment in the given organ. For leaves, this gain comes through increased light capture and photosynthesis; for roots, through improved water uptake that relieves a limitation on photosynthesis elsewhere in the plant. In principle, roots can also improve nutrient acquisition and uptake but that is currently outside the scope of DAESIM2-Plant. The marginal cost determines how rapidly a unit of carbon would be lost, given the organ’s typical turnover rate. Marginal gain–cost ratios are normalized across pools so that allocation fractions sum to unity, and negative economic gains are set to zero, so that a pool is not allocated carbon when doing so would reduce net carbon profit.

At each time step and for each plant carbon pool, *C*_*k*_, the marginal carbon gain is evaluated with:

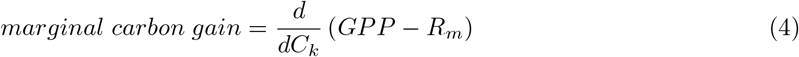

And the marginal carbon cost from allocating to that pool:

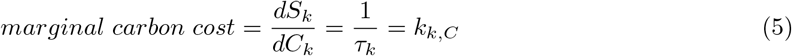

Where *τ*_*k*_ equals the mean lifespan of pool *k* (days) and *k*_*k,C*_ is its turnover rate (days-1). This ensures that carbon invested in short-lived pools is a costlier investment than carbon invested in long-lived pools. In this way, the marginal gain term is calculated directly from the coupled photosynthesis, stomatal conductance, and plant hydraulics modules described above. Any trait that alters those processes, for example, higher photosynthetic capacity (*V*_*cmax*_) or greater root hydraulic conductance changes the marginal gain of investing in a given organ, and therefore changes the allocation strategy that emerges, without any explicit rule linking that trait to allocation. Equally, the same trait can produce different allocation outcomes in different environments, because the marginal gain depends on prevailing conditions such as soil moisture, radiation, or temperature. In this way, the optimal allocation framework provides a link between plant traits, environmental conditions and growth.

The allocation coefficients are determined numerically. In only a limited set of cases is there an analytical solution. For example, (Potkay et al., 2021) showed that under certain assumptions this is the case, while Thornley and Parsons (2014) showed that analytical solutions to teleonomic models are possible under assumption of exponential growth. The gradient ascent method applied here uses only local slope information, and so is not guaranteed to find the single best allocation within a given step, only to move in approximately the right direction. Potkay et al. (2021) showed this approximation to be a good match against a numerical solution. Utilising a numerical approach has the added benefit of easier ongoing model development and testing of alternative process hypotheses, such that analytical solutions do not need to be derived each time.

#### 3.1.5 Plant Development and Phenology

Plant development is governed by the accumulation of heat units, or thermal time, which determines the timing of transitions between developmental phases (e.g. germination, vegetative growth, anthesis, grain filling, maturity). Thermal time was chosen because temperature-based development timing is well supported empirically, and GDD requirements are widely reported for many crops and cultivars (e.g. Celestina et al., 2023), allowing the model to be readily parameterized from existing data.

No single method for calculating thermal time is standard, and different methods are not directly interchangeable (McMaster and Wilhelm, 1997). DAESIM2-Plant therefore offers two linear methods, which are simple and perform well at intermediate temperatures (matching how most published GDD requirements are derived), and a non-linear method based on a beta distribution function with a thermal optimum, which better represents the underlying biological response at temperature extremes (Bonhomme, 2000) and is preferable for future climate scenarios or population-level simulation. The non-linear method requires hourly temperature, yet often only daily data are available, so a synthetic diurnal cycle is generated from daily minimum and maximum temperatures using the WAVE model of Bal et al. (2023).

Vernalization is included for cultivars in which it meaningfully affects flowering time (e.g. winter wheat), with linear (Wang and Engel, 1998), non-linear sigmoidal (Streck et al., 2003), and APSIM-Wheat-style response functions provided, again to allow the model to be matched to whatever cultivar-specific parameters are available.

Developmental phase is the primary link between development and the rest of the model: each phase sets its own GDD requirement, organ-specific turnover rates, and (for stem and grain) fixed allocation coefficients, and triggers processes such as spike growth (Section [Grain Production]) and the maturity-phase downregulation of photosynthetic and hydraulic capacity. Leaf and root allocation within each phase continues to respond dynamically via the optimal allocation framework described above.

#### 3.1.6 Wheat Grain Production

Grain production in DAESIM2-Plant is represented using a source–sink approach. The sink, represented by the potential number and size of grains the crop can produce, is determined using the concept of the critical period, a defined developmental window around anthesis during which floret survival and spike growth determine the final grain number. This is based on extensive eco-physiological evidence, reviewed by Fischer et al. (2024) and Pretini et al. (2021), showing that grain number in wheat is set by the assimilate supply to the developing spike during this period rather than by conditions during grain filling itself, and that it correlates strongly with spike dry weight at anthesis (Fischer and Stockman, 1980; Terrile et al., 2017). The source is determined by the flow of new assimilates from ongoing photosynthesis together with the re-translocation of assimilates stored in the stem, reflecting evidence that stem water-soluble carbohydrates act as an important buffer for grain filling in wheat, particularly under stress. Stem reserves typically contribute up to 20% of final grain weight under favourable conditions, but can compensate for 50-70% of grain weight when photosynthesis is limited by terminal drought or heat stress (Blum, 1998).

Target grain number and weight. The potential grain density, 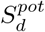, is calculated from the spike dry weight at anthesis, *SDW*_*a*_, and a genotype-specific fruiting efficiency, *FE*, which quantifies the reproductive efficiency of the spike in setting grains per unit of spike growth:

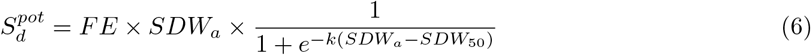

The sigmoid term, with parameters *k* and *SDW*_50_, modulates the relationship at low and high extremes of *SDW*_*a*_, consistent with the near-linear relationship observed between spike dry weight and grain number across most of the observed range, but with a saturating response as *SDW*_*a*_ becomes very large and a non-linear decline as it approaches zero (Fischer et al., 2024; Pretini et al., 2021), which is perhaps useful when representing a population of plants. Because DAESIM2-Plant does not represent the spike as an organ distinct from the stem, *SDW*_*a*_ is inferred as the increase in stem biomass during the spike-formation development phase, prior to anthesis, meaning that environmental or trait-based effects on photosynthesis and developmental timing during this phase propagate directly to potential grain number. Potential grain weight, 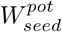 , is then obtained by multiplying 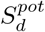 by a fixed mean individual grain weight, *GW* , since variation in grain weight explains comparatively little of the variation in wheat yield relative to grain number (Peltonen-Sainio et al., 2007).

Fruiting efficiency, *k*, and *SDW*_50_ are treated as relatively conservative parameters (typical ranges FE=80-210 *grains g d*.*wt*^−1^; k=0.01-0.03; *SDW*_50_=80–150 *gd*.*wtm*^−2^, consistent with observed ranges of *SDW*_*a*_ (100-250 g d.wt m-2), grain number (roughly 2000-19000 grains m-2 across spring and winter wheat), and individual grain weight (28-48 mg) reported across multi-site trials (Peltonen-Sainio et al., 2007; Terrile et al., 2017). Of these, *FE*, is expected to vary most strongly by genotype and is therefore the primary parameter distinguishing cultivars in the grain production module, while *k* and *SDW*_50_ are held approximately constant.

Grain filling. Once the potential grain pool size is set, whether it is achieved depends on assimilate supply during grain filling. New assimilates are allocated to the grain pool via a fixed allocation coefficient once the developmental phase transitions to grain filling. Stem remobilization is represented as a Michaelis–Menten function of the concentration of remobilizable carbon in the stem (Supplement Equation X), reflecting the assumption that remobilization is limited by the rate of phloem loading rather than by the size of the stem reserve pool alone, based on phloem-loading kinetics described for trees (De Schepper and Steppe, 2010; Trugman et al., 2018). To avoid the need to explicitly track non-structural carbohydrate pools separately from structural stem biomass, we use the ratio of stem to leaf carbon, *C*_*stem*_ : *C*_*leaf*_ , as a proxy for the concentration of remobilizable stem carbon, since this ratio is elevated during spike formation (when remobilizable reserves are high) and declines as structural growth continues. Both sources continue to supply the grain pool until either the grain-filling phase ends or the potential grain biomass, 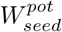 , is reached, whichever occurs first. This way, realized yield is jointly constrained by the sink established during the critical period and the source available during filling.

#### 3.1.7 Coupling Carbon Allocation Across Organs

The allocation coefficients to the four organ pools including leaves (*u*_*L*_), roots (*u*_*R*_), stem (*u*_*S*_), and grain (*u*_*G*_), must sum to one at every time step. In the current implementation, only *u*_*L*_ and *u*_*R*_ follow the optimal trajectory principle described above, while *u*_*S*_ assumes fixed minimum value that is phase-specific, and *u*_*G*_ follows the critical-period/grain-filling rule described in Grain Production, taking a fixed maximal value while the actual grain pool remains below its potential size, 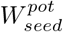 , and zero otherwise. The optimal fractions, denoted 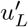 and 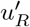 (with 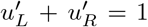), are therefore not applied directly but get scaled by the other terms as follows:

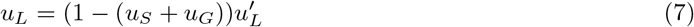

and

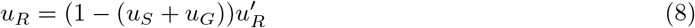

This preserves the relative split between leaves and roots that the optimality principle selects, while ensuring the four coefficients sum to one at every time step. A practical consequence is that leaf and root allocation is not optimal in isolation: it is the optimal division of whatever carbon remains once stem growth or active grain filling has taken its fixed or threshold-driven share. A phase with high fixed stem allocation, or a period of active grain filling, therefore directly constrains how much carbon the optimal scheme can invest in leaves or roots, independent of how favourable conditions for photosynthesis or water uptake might otherwise be, ensuring sensible coupling between developmental structure of the model and optimal allocation strategy.

### 3.2 Model Experiments

### 3.3 Idealised Sensitivity Experiments

To isolate how the optimal carbon allocation scheme responds to plant state and environmental conditions, we conducted three sets of idealised experiments. For these experiments the model’s instantaneous allocation response was diagnosed at prescribed, fixed combinations of leaf biomass, root biomass, and soil moisture. This provides insight into the allocation dynamics independent of a full developmental trajectory over a growing season. These experiments used a three-layer canopy and a single-layer soil for simplicity, with the same parameterization as the field-site simulations (Section 2.3).

In the first experiment, leaf biomass was varied between 0 and 500 g d.wt m-2, while holding root biomass fixed at 100 g d.wt m-2. Second, root biomass was varied over the same range while holding leaf biomass fixed at 100 g d.wt m-2. Both were repeated across five volumetric soil moisture levels ranging from very dry to near saturation (0.25-0.45 m3 m-3). This isolates the direct effect of increasing investment in one organ from the state of the other organ and from prevailing soil moisture, diagnosing how net carbon profit (GPP - Rm), transpiration rate, and the resulting leaf and root allocation coefficients respond as each pool is built up under a given environmental condition.

In the third experiment, volumetric soil moisture was varied continuously (0.20–0.50 m3 m-3) at four fixed root:leaf biomass ratios (0.05, 0.2, 0.5, and 1), diagnosing the same four outputs. Because the root:leaf ratio reflects the plant’s structural investment strategy while soil moisture is a purely environmental driver, this isolates how the same environmental gradient produces different allocation outcomes depending on the plant’s current structural balance between above- and below-ground tissue.

### 3.4 Idealised Growing Season Experiments

To understand how the model responds to sustained water constraints over a growing season, we ran the model over a growing season under a range of fixed soil moisture levels and daily varying meteorological forcing. This provided an idealised but plausible set of simulations to understand how instantaneous allocation evolves in combination with crop development, carbon uptake, and grain yield.

The model was run at a representative location in the wheat belt of south-western Australia, which is situated in a Mediterranean climate zone. Five simulations were run under five relative soil wetness levels, expressed as the fraction of the plant-available water capacity (PAWC). The PAWC which represents the relative difference between the drained upper limit and the crop lower limit. The chosen PAWC levels were 0.2, 0.3, 0.4, 0.5, and 0.6. The lowest PAWC level represents a dry soil close to the crop lower limit while the uppermost PAWC level represents a moderately wet soil with ample moisture available for crop water use. With this experimental design, we could analyse the effect of water availability through a complete growing season, from sowing to maturity, in the presence of typical variation in meteorology and crop development. Meteorological forcing was retrieved from the ANUClimate data collection version 2 (Michael et al., 2021) for the selected location. The configuration of the plant model was the same as above, with a three layer canopy and single layer soil for simplicity.

## 4 Results

### 4.1 Sensitivity to Leaf and Root Biomass

At constant root biomass, net carbon profit and transpiration rate both increase with leaf biomass following a positive, curvilinear relationship. This relationship saturates at a soil moisture dependent asymptote (Fig. 1a,b), reflecting two compounding limits. First, as canopy leaf area approaches its maximum, further leaf growth provides little additional light interception. Second, with a fixed root biomass and soil moisture, water uptake increasingly caps transpiration, and hence photosynthesis, that additional leaf area can support. At any given leaf biomass, both net carbon profit and transpiration are higher under wetter soil conditions. Under the driest soil moisture level, both remain low and are effectively unresponsive to further leaf growth, since water availability limits gas exchange well before maximum LAI is approached.

**Figure 1:**
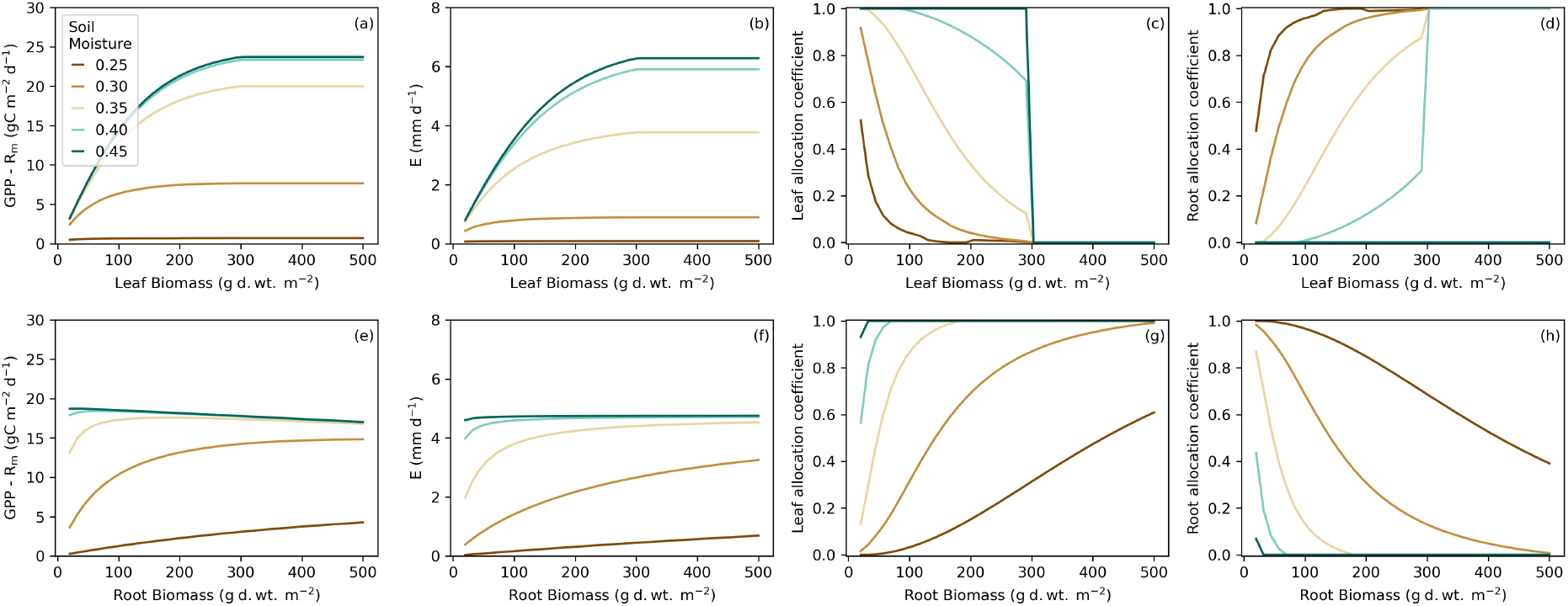
Simulations showing the sensitivity of DAESIM2 to the leaf biomass and root biomass across five soil moisture levels. Figures show net carbon profit (GPP - Rm), transpiration rate (E), leaf allocation coefficient and root allocation coefficient.

This saturating response has a direct consequence for allocation. As leaf biomass increases, the marginal gain from further leaf investment declines, driving a corresponding decline in the leaf allocation coefficient and a complementary increase in the root allocation coefficient (Fig. 1c,d). This represents a diminishing return on carbon invested in leaves as leaf biomass increases while root biomass and soil moisture remain fixed. Under drier soil conditions, this diminishing return sets in earlier and is more pronounced, so the model allocates relatively more new carbon to roots than it would under wetter conditions at the same leaf biomass, consistent with the well-documented plasticity of biomass partitioning under water stress (Vandoorne et al., 2012).

At constant leaf biomass, the response to increasing root biomass differs qualitatively with soil moisture (Fig. 1e-h). Under wetter conditions, transpiration is essentially insensitive to root biomass (Fig. 1f), indicating that root water uptake capacity is not limiting. In this case, net carbon profit declines approximately linearly with root biomass (Fig. 1e) due to the additional maintenance respiration cost of greater root biomass. Under drier conditions, a positive curvilinear relationship develops between root biomass and both transpiration and net carbon profit, as additional roots relieve a water-uptake limitations on net carbon profit. This soil moisture dependent transition, whereby root growth incurs only a respiratory cost under wet conditions versus a beneficial investment under dry conditions, is produced endogenously, without a prescribed water-stress response function on carbon allocation itself.

### 4.2 Sensitivity to Soil Moisture

Figure 2 shows these dynamics across a continuous soil moisture gradient at four fixed root:leaf biomass ratios. Net carbon profit and transpiration increase with soil moisture up to a plateau (Fig. 2a,b), beyond which further increases provide no additional benefit. This point indicates where soil moisture ceases to limit gas exchange, which is governed by the underlying soil-plant-atmosphere continuum. Thus, this point will shift depending on soil type and plant hydraulic properties, including the parameters governing the relationship between leaf water potential and stomatal conductance in the Tuzet model. With increasing soil moisture, leaf allocation increases and root allocation correspondingly decreases (Fig. 2c,d). Under moist conditions, the model favours leaf investment to capture more light, since water is no longer scarce. Under dry conditions it favours roots to relieve the water constraint. The root:leaf ratio changes the soil moisture level at which the model shifts new investment from roots back to leaves, or vice versa. A plant with a higher root:leaf ratio can tolerate drier soil before its roots cannot fully support water loss via transpiration and it needs to invest new assimilates there, so the shift toward leaf allocation happens at a drier soil moisture. A plant with fewer roots relative to leaves needs wetter soil before the same shift occurs, since it would have less root capacity to draw on. This shows that allocation depends not just on soil moisture but on how much root water uptake capacity the plant already has relative to its demand via transpiration.

**Figure 2:**
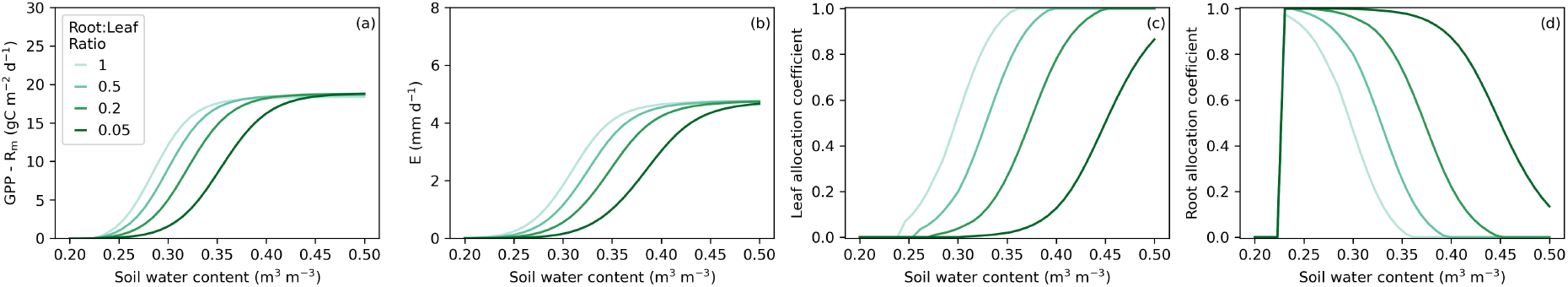
Simulations showing the sensitivity of DAESIM2 to soil moisture across four root:leaf biomass ratios. Figures show net carbon profit (GPP - Rm), transpiration rate (E), leaf allocation coefficient and root allocation coefficient.

### 4.3 Growing Season Sensitivity to Plant Available Soil Wetness

Simulating a full growing season under a gradient of relative PAWC levels allows us to test the model response to sustained water constraints. For these simulations, soil moisture was held constant throughout the season at each of five PAWC levels, isolating the effect of overall water availability from any dynamic soil-drying process. This allows us to evaluate whether the instantaneous allocation response demonstrated in the idealized experiments above translates into realistic patterns of canopy development, carbon uptake, biomass partitioning, and grain yield across a full season.

Canopy development (LAI) and carbon uptake (GPP) show a strongly saturating response to PAWC (Fig. 3a,b). The three wettest treatments (0.4–0.6) all reach the maximum LAI and track closely in both LAI and GPP trajectories, indicating that once PAWC exceeds a threshold (dependent upon traits and environment), further increases in water availability provide comparatively little benefit to canopy growth. The 0.3 treatment develops more slowly and peaks at a substantially lower LAI and GPP. The driest treatment (0.2) fails to develop a functional canopy at all, as LAI peaks at approximately 1.2 m2 m-2 and GPP remains low (less than 3 g C m-2 d-1) for the entire season. This non-linear, threshold-like pattern, whereby three treatments perform similarly well, one intermediate, and one representing effective crop failure, illustrates that PAWC does not constrain growth proportionally across its full range, but instead becomes acutely limiting only below some critical level.

**Figure 3:**
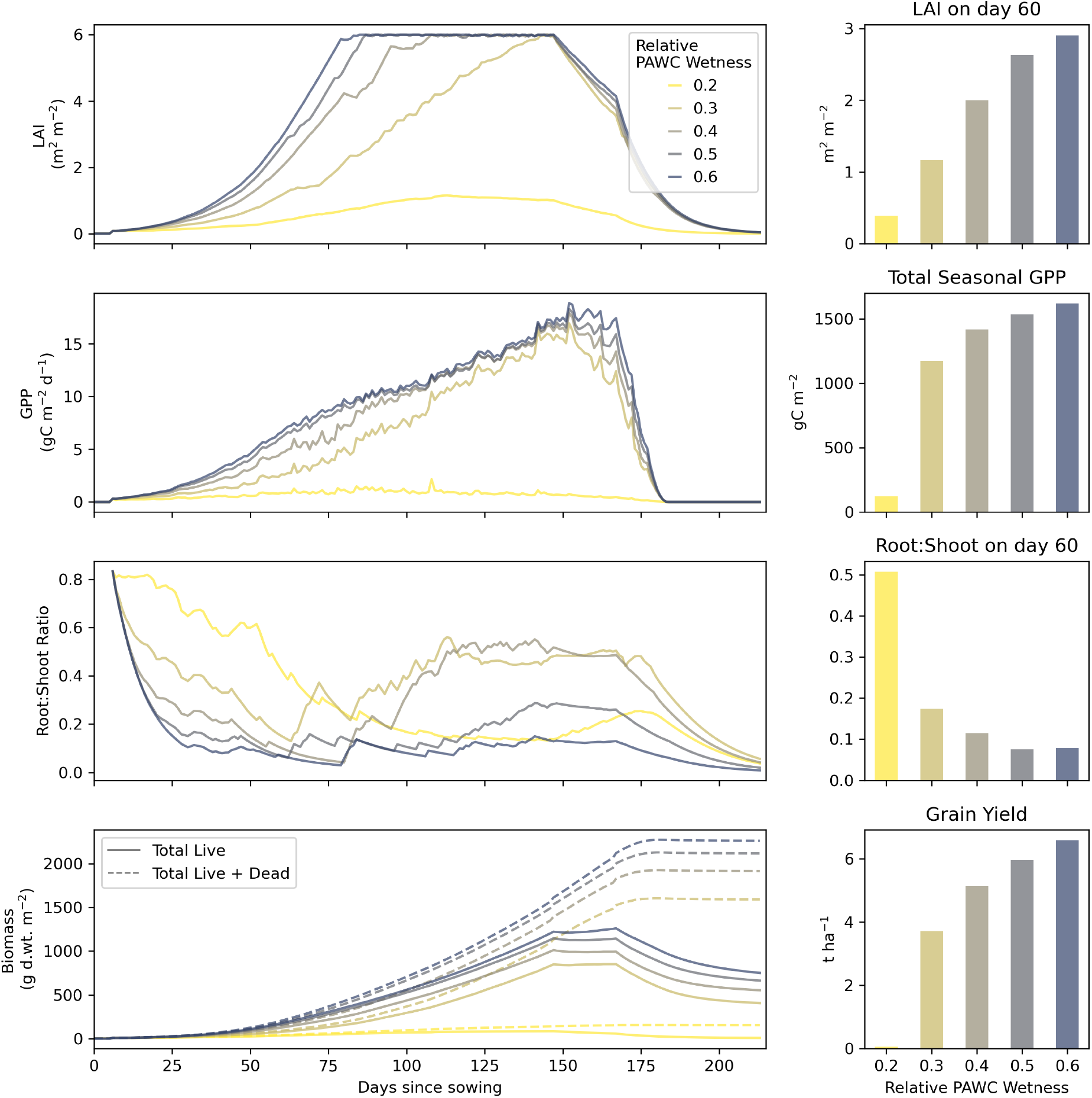
Idealised model simulations of wheat over a growing season across five constant plant-available water capacity (PAWC) levels in a Mediterranean climate. The root:shoot ratio is the ratio of total root biomass relative to combined leaf and stem biomass. Total live biomass includes the leaf, root and stem pools and excludes the grain pool.

Under the driest treatment, the root:shoot ratio remains elevated throughout the season (typically 0.3–0.65) rather than declining to the low values reached under wetter conditions (Fig. 3c). This is summarized concisely by root:shoot ratio on day 60 since sowing (Fig. 3g), which falls from approximately 0.5 under the driest PAWC to approximately 0.08 at the wettest PAWC, with most of this decline occurring between PAWC levels 0.2 and 0.4 and comparatively little further change beyond that point. This mirrors the pattern seen in the earlier idealised experiments (Figs. 1, 2): as water becomes more limiting, the model shifts new carbon investment toward roots, and this shift saturates once soil moisture is no longer the dominant constraint on growth.

Total biomass accumulation, including both living and dead/senesced carbon, (Fig. 3d) reflects a similar saturating pattern seen in canopy development and carbon uptake. With drier PAWC levels, by the end of the growing season the biomass accumulation plateaus at progressively lower totals. Under the wettest PAWC, total biomass accumulation reaches about 2200 g d.wt m-2, the second driest PAWC level reaches about 1600 g d.wt m-2 while the driest treatment is below 200 g d.wt m-2. As expected, this closely mirrors the total seasonal GPP (Fig. 3f). The total live biomass (leaves, roots, stems) shows a similar relative difference across PAWC treatments albeit with a slightly different growth trajectory late in the growing season. Total live biomass peaks around day 145 since sowing, coinciding with the start of the anthesis developmental phase. Grain filling starts on day 165 since sowing, which begins to draw down the stem pool as non-structural carbohydrates are re-translocated to the grain. This also coincides with higher turnover rates of the leaves, roots and stems which is further enhanced during the senescence developmental phase as determined by the phase-specific turnover parameters.

This behaviour offers a way to interpret the model in relation to the well-documented association between soil texture and root:shoot allocation (Ding et al., 2025), whereby the low water-holding capacity of sandy soils leads to additional carbon investment in roots to sustain water uptake. Because soil moisture is prescribed in DAESIM2-Plant rather than simulated dynamically, we cannot directly represent the two-way interactions between soil moisture and plant water uptake. However, soil texture is a primary determinant of PAWC (Gladish et al., 2021), and the model’s PAWC gradient can be interpreted as a proxy for a gradient in effective soil texture, from sand-like (low PAWC, 0.2) to loam- or clay-like (higher PAWC, 0.5–0.6) conditions. Under this interpretation, the pronounced increase in root:shoot ratio at low relative PAWC (Fig. 3c) is qualitatively consistent with the empirical expectation that sandy, low-water-holding-capacity soils drive greater relative investment in root biomass. This is qualitative, rather than quantitative, correspondence because water uptake and soil drying are not dynamically coupled to a specified soil texture in this model configuration, we cannot say to what degree a given sand fraction would translate into a given PAWC value, only that the direction and general shape of the allocation response to reduced water availability is consistent with what is observed for sandy soils in the field.

One aspect of these results also warrants further comment. The root:shoot ratio trajectories under the driest treatments are visibly noisier than those under wetter conditions (Fig. 3c). This is likely caused by two compounding features of the model. First, allocation is solved instantaneously at each time step (daily) with no temporal smoothing or weighting back in time. Thus, day-to-day variability in drivers of net carbon profit will impact allocation dynamics, such as air temperature, vapour pressure deficit and incoming shortwave radiation. Second, stomatal conductance and canopy photosynthesis become more sensitive to day-to-day variations in vapour pressure deficit under water-limited conditions, since leaf water potential runs closer to the threshold at which stomatal down-regulation occurs. Moderate fluctuations in daily VPD can therefore drive relatively large swings in net carbon profit. Because net carbon profit is itself small in absolute terms when the soil is dry, these fluctuations may be further amplified in the marginal gain-cost ratio that determines the allocation fractions, producing more day-to-day variations. This is not an inherently problematic aspect of model behaviour. However, this could be addressed in future by moving from an instantaneous approach to a temporally weighted approach of determining the marginal carbon gains and marginal carbon costs. Similar temporal weighting was applied in Caldararu et al. (2020) in their whole-plant optimality approach to modeling leaf nitrogen changes.

Crop yield (Fig. 3h) mirrors the threshold-like response evident in carbon uptake. The yield is near zero (less than 0.1 t ha-1) under the driest treatment, consistent with effective crop failure due to insufficient soil water. The second driest treatment shows a relatively large increase in crop yield to 3.7 t ha-1, followed by progressively smaller increases in crop yield as PAWC increases, up to 6.6 t ha-1 in the wettest treatment. This follows directly from the fact that crop yield in this model is constrained by the same assimilate supply that determines vegetative growth during both the critical period, which coincides with spike formation and stem assimilate storage, and the grain filling period via new assimilates. It is important to note that these idealised simulations hold the soil moisture at a fixed level across the growing season. In reality, the timing of a dry spell relative to these two periods is likely to be important. A dry spell during spike formation may limit the potential grain number (sink strength) and storage of non-structural carbohydrates in the stem. Crop yield is likely to be reduced even if water limitation is alleviated during grain filling and direct assimilate supply is adequate. Alternatively, favourably wet conditions during spike formation followed by a late season dry spell could produce strong sink strength and ample stem reserves, yet still restrict crop yield if new assimilate supply during grain filling is low. The temporal dynamics of soil moisture, not just its seasonal average, are likely to be an important control on yield outcomes in this model. The source-sink structure of the grain production module is well placed to be able to test such scenarios in future work.

## 5 Discussion

### 5.1 The Optimal Path Assumption

There is a long history of so-called teleonomic models describing carbon allocation and these share a common structure. Each defines a target state, typically a particular root:shoot ratio or resource stoichiometry, and constructs plant dynamics to seek and maintain it (Thornley, 1972a,b, 1995; White, 1937; Brouwer, 1962). These have many names: Functional equilibrium, functional balance, balanced growth, optimal partitioning theory, and eco-evolutionary optimality. The optimal path approach used here, following Buckley and Roberts (2006), Caldararu et al. (2020) and Potkay et al. (2021), shares this lineage but departs from it in one specific respect. The objective is fixed, whereby carbon is allocated to maximize potential net carbon profit, but the solution is not. The marginal gain and cost are recomputed at every time step from the current plant state and environmental conditions, so the allocation that satisfies the objective shifts continuously rather than converging on a predetermined ratio. This distinguishes an optimal path from an optimal allometry: solving the same objective repeatedly against changing conditions, rather than solving once for a fixed destination. With this approach, the allocation scheme carries no memory of prior states and does not perform forecasting of future conditions.

This distinction addresses one contentious issue with teleonomic allocation models (Robinson, 2022). With no encoded destination, nothing in the model requires convergence toward a fixed ratio after a perturbation, nor tightly balanced root and shoot growth rates. Our idealized experiments show strongly unbalanced allocation as the norm rather than an exception to be explained away. Apparent recovery toward a prior allocation state after a perturbation is better read as the same physiological and environmental drivers reconstructing a similar marginal-gain landscape, not as goal-directed correction. Where leaf and root allocation are coupled, that coupling is not asserted a priori. Instead, it emerges because both organs’ marginal returns are evaluated against a shared carbon and water budget, linked through the soil-plant-atmosphere hydraulic pathway.

This distinction is important and provides a plausible description of how carbon allocation behaves, although not with the mechanistic rigour of other approaches that explicitly represent phloem transport and local utilization (e.g. transport-resistance approaches). As within the eco-evolutionary optimality tradition more broadly, allocation is best understood as continuously re-evaluated against a changing environment, with optimality approached rather than ever fully achieved (Harrison et al., 2021). The framework remains phenomenological as it does not specify the physiological mechanisms by which a plant would compare marginal returns across distance organs, via sugar or hormone signaling, for example. It describes how allocation shifts with trait and environment, not why, mechanistically, a plant would execute that shift. In any case, the proximate mechanism is difficult to test directly against experimental data, though the models emergent outputs including allocation coefficients, biomass, and yield, remain testable. The choice of fitness proxy also remains a subjective modeling decision. What the approach offers is a parsimonious, dynamically responsive alternative to fixed or empirically prescribed allocation rules, generating context-dependent behaviour rather than assuming it.

### 5.2 The Optimality Assumption for Crops

As Fernie et al. (2020) note, the adaptive strategies plants evolved for fluctuating natural environments are not always required in agricultural systems, and can even constrain yield. The assumption of an optimal allocation warrants particular scrutiny for crops, since modern cultivars are products of intensive artificial selection for agronomic outcomes, particularly yield and harvest index, rather than natural selection for fitness in a competitive, uncultivated environment.

One clear example is the harvest index. The semi-dwarfing genes underpinning the Green Revolution in wheat and rice (e.g. Rht-B1b, Rht-D1b; sd-1) reduce sensitivity to gibberellin hormones, suppressing stem elongation and redirecting assimilates to the spike rather than competing stem growth (Hedden, 2003; Youssefian et al., 1992). Taller stems improve light competition in a wild stand but are selected against for a dense, fertilized monoculture, an outcome is difficult to characterize as fitness-optimal. This is one justification for keeping allocation to the stem as phase-specific coefficients in DAESIM2-Plant, rather than treating it as optimal. Stem allocation follows fixed, phase-specific coefficients, while grain allocation follows a critical-period and threshold rule. Only the leaf:root split follows the optimal path hypothesis. Root architecture has, by contrast, been a comparatively minor and largely unconscious breeding target relative to harvest index (Zhu et al., 2019; Reynolds et al., 2021), and the light-versus-water trade-off our optimality principle acts on remains directly relevant to yield under variable field conditions.

We would still distinguish the optimality of the resulting strategy from the plasticity of the underlying response. The evidence motivating this approach is not that crops allocate carbon in a way that is demonstrably optimal, but that allocation responds dynamically and predictably to resource limitation and plant traits, as documented in modern cultivars (Vandoorne et al., 2012; Chandregowda et al., 2023; Bacher et al., 2022). Eco-evolutionary optimality is used here as a parsimonious means of generating this behaviour without prescribing it directly. Furthermore, other features of the model are prescribed as fixed genotype parameters, such as maximum canopy height, maximum leaf area index, and maximum rooting depth. These impose hard genotypic constraints on canopy and root system development, but the optimality scheme adjusts allocation natively to accommodate them rather than requiring this to be built into the allocation module itself. Whether breeding has shifted the specific trade-offs the model resolves is an empirical question, testable by comparing model behaviour against modern, historical, or wild cultivars. The value of the approach lies in generating realistic, emergent behaviour from few well-justified assumptions rather than many prescribed rules, and less in whether the plant is truly optimal.

### 5.3 Could optimality be extended to the stem and grain?

One limitation of this model concerns the stem and grain pools, which currently do not feed back into plant function. The stem acts largely as a passive carbon reservoir, as it does not contribute to photosynthesis, transpiration, or hydraulic conductance; although it can be used to supply carbon during grain production via retranslocation. Similarly, the grain does not contribute plant function. This reflects a broader open question this model does not resolve: how much of carbon allocation is genetically constrained versus an emergent, plastic response to conditions. For leaves and roots, we address this with genotype-specific trait limits — maximum LAI and maximum rooting depth — within which allocation is otherwise governed by the optimality scheme; once a limit is reached, marginal gain for that organ falls and carbon is redirected elsewhere (Fig. 1c). We do not yet apply an equivalent structure to stem or grain.

A natural extension would be to give the stem an explicit function. For instance, hydraulic resistance along the transport pathway, structural support, and/or storage capacity, so that it enters the same marginal-gain calculations as leaves and roots, but bounded by a genetically determined limit, analogous to maximum LAI. This would let the optimality scheme govern stem investment up to that limit, rather than treating it as an unconstrained sink once formed, while preserving the reality that breeding has directly targeted the eventual balance between stem and grain. This would also reduce the number of free parameters needed to assign as part of the phase-specific stem allocation. The same logic could, in principle, extend to grain, though its distinct source-sink dynamics during the critical period would require separate treatment. This as an enticing but largely untested direction. Implementing a clear functional role for the stem is likely to be necessary if extending DAESIM2-Plant to non-herbaceous species in future. In trees, for example, hydraulic resistance along the stem becomes an increasingly important constraint as the hydraulic path length increases dramatically with tree height (Potkay et al., 2021), scaling with stem cross-sectional area via the Hagen–Poiseuille relationship.

### 5.4 Soil Texture, Nutrient Availability, and Root:Shoot Allocation

Our interpretation of the PAWC sensitivity experiment in relation to soil texture is supported by more detailed evidence on the relationship between soil texture and root:shoot allocation in cereals. Poeplau and Kätterer (2017) found that root:shoot ratios in spring barley decreased significantly with increasing clay content (or, equivalently, decreasing sand content), a pattern also reported across several other crop types (Ding et al., 2025). The mechanism proposed is that reduced root–soil contact in sandy soils — rather than a light, *CO*_2_, or nutrient limitation — drives greater carbon investment belowground to maintain hydraulic conductance at the root-soil interface (Ding et al., 2025). Notably, Poeplau and Kätterer (2017) found this texture effect on root:shoot ratio to be independent of long-term fertilizer history, even though total shoot and root biomass were both higher in clay-rich soils, potentially reflecting better nutrient access rather than a shift in allocation strategy per se.

This is a useful point of contrast with nutrient-driven allocation shifts reported elsewhere. Hansson et al. (1987) found that nitrogen fertilization in barley increased total biomass while decreasing root:shoot ratio, relative to unfertilized controls with lower total biomass and a higher root:shoot ratio — consistent with a functional-equilibrium-type adjustment of allocation toward whichever resource (in this case, nitrogen) is most limiting. Taken together, these two lines of evidence suggest that root:shoot allocation in cereals responds to soil texture and nutrient availability through at least partially distinct mechanisms — one hydraulic (root-soil contact and water uptake efficiency), the other resource-economic (nitrogen supply relative to demand) — both of which are qualitatively consistent with the kind of trait-by-environment interplay this paper’s optimality framework aims to capture. Since DAESIM2-Plant does not currently represent nutrient limitation (Section 2, Methods scope statement), the model can speak only to the hydraulic mechanism; extending the optimal allocation framework to include a nitrogen-limited marginal gain term — for example, following the nitrogen-allocation optimality approach of Franklin (2007) — would be a natural way to test whether both mechanisms can be captured within the same theoretical structure, rather than requiring separate, mechanism-specific allocation rules.

### 5.5 Towards Evaluation Against Field Data

The results presented here are drawn entirely from idealized sensitivity experiments, designed to isolate and diagnose the behaviour of the optimal allocation scheme under controlled combinations of plant state and environmental conditions. This is a deliberate first step in describing a new model and its theoretical basis, but leaves open the question of how well DAESIM2-Plant reproduces observed growth, allocation, and yield outcomes under real field conditions. A clear next step is to evaluate the model against experimental and field trial data, including datasets that capture drought stress and other water-limited conditions, since these are precisely the conditions under which the optimal allocation scheme is expected to produce the most distinctive, trait- and environment-dependent behaviour. Such evaluation would also provide a direct test of the qualitative claims made throughout this paper against quantitative outcomes rather than idealized theoretical expectations alone.

## 6 Conclusions

The DAESIM2-Plant model represents a significant step forward in crop simulation, offering a flexible and mechanistically grounded framework for understanding plant growth and resource allocation across environmental conditions. By moving beyond empirical coefficients and utilising trait-based, mechanistic formulations, in combination with an eco-evolutionary optimality principle for carbon allocation, the model may enhance our ability to predict crop responses to environmental variability and supports the development of resilient agricultural systems.

Meeting the dual demands of expanding productivity and building resilience requires tools that generalize beyond the specific conditions under which they were developed. DAESIM2-Plant works toward this by constraining growth to known physiological limits, rather than requiring exhaustive calibration of genotypes under specific environmental conditions. This is intended to provide a more generalizable model that can be applied beyond the conditions used for its development, including novel or under-sampled combinations of climate, soil, and management. Future work will focus upon evaluation against observations, and extensions of this framework to other major crop types, including perennial grasses, C4 species, legumes, and woody perennials, would allow the same underlying principles to be applied across a broader range of agricultural and natural systems.

## 7 Acknowledgements

JB was supported by an ARC Linkage Project LP190101060 to The Australian National University Project title Integrated Farm Modelling to Improve Resilience and Sustainable Prosperity. AN and JB were supported by the Commonwealth Department of Environment and Energy - Ext led by CSIRO - National Environmental Science Programme 2 (NESP) Climate Systems Hub - NESP 2 Activity Schedule 3.

