## Supplementary Material for "Eco-Evolutionary Optimal Carbon Allocation in a Mechanistic Crop Growth Model: Theory and Application to Wheat"

September 21, 2026

The Dynamic Agro-Ecosystem SIMulator 2 is a plant growth and development model (DAESIM2-Plant) that simulates key physiological, structural, and developmental processes that govern plant carbon, water and potential yield dynamics over time. The model integrates physiological processes such as photosynthesis, stomatal conductance, and water uptake with developmental dynamics, including growth stages, carbon allocation to various organs, biomass turnover and grain production. Carbon allocation is dynamically influenced by plant physiology, developmental stages and environmental conditions, ensuring realistic representation of plant growth trajectories. Grain production is crop type specific, with a focus on so-called critical periods, and links to the rest of the structural and functional plant processes. By coupling plant physiology with development and structure, this model provides a comprehensive framework for understanding and predicting plant responses to environmental factors and management practices.

#### 1 Plant Structure

The model explicitly represents four primary live plant organs: leaf, stem, root, and seed (grain), each contributing uniquely to resource acquisition and overall plant function. Plant structure and physiology are intricately linked, as the composition and arrangement of organs determine the efficiency of certain processes. For instance, the structure and orientation of leaves and stems regulate light interception, driving photosynthesis. Similarly, root structure and hydraulic properties govern water uptake from the soil, influencing transpiration rate and water use efficiency. These interactions underscore the importance of organ-specific contributions to whole-plant performance. The model is designed to represent a population of plants with homogenous traits and structure. The model assumes a horizontally uniform canopy albeit with some important features, such as shading within the canopy and foliage clumping effects.

#### 2 Plant Development

Plant development in the model simulates the progression of a plant through distinct developmental phases, which are defined by temperature through exposure to heat (growing degree days, GDD) and cold (vernalization). The model is highly flexible, allowing for an arbitrary number of developmental phases, as long as phase-specific parameters are assigned. For each phase, the model requires values for GDD requirements (the amount of temperature exposure needed to progress), vernalization requirements (if relevant), carbon allocation coefficients (which determine how much carbon is directed to different parts of the plant), and turnover rates (biomass loss over time). The turnover rates govern the conversion of live biomass to dead litter biomass. Within each phase, the carbon allocation coefficients must sum to equal 1. These shifts in carbon allocation coefficients and turnover rates reflect the expected changes in the growth and development of plant organs as the plant progresses through its life cycle.

In the generic crop configuration, the model is parameterized with five developmental phases: germination, vegetative growth, anthesis (flowering), fruiting (grain production) and maturity. Each phase has unique characteristics. Germination is the initial phase, requiring relatively few GDD for the seed to germinate and emerge. During this phase, carbon is almost entirely allocated to the roots to establish the seedling. Vegetative growth follows germination, where the transition point represents emergence.

During the vegetative phase, the GDD requirement is typically high and the plant focuses on expanding its leaf and root biomass to capture resources. Anthesis marks the transition to reproduction, where the plant begins to allocate carbon toward flowering and seed formation. Fruiting is the final phase, where a large proportion of carbon is allocated to seeds. Following seed production an annual plant has completed its reproductive cycle and has reached maturity and begins to senesce, with turnover rates becoming higher and physiological function declines.

#### 2.1 Thermal Time

The rate of development in annual crops is closely tied to temperature, which reflects the available energy for plant processes. The principle is that certain temperatures are required for the plant to progress through its life cycle. The concept of heat units or thermal time is widely used to quantify, describe and predict the developmental rate based on temperature. Mathematically, thermal time is represented as the integral of a temperature curve over time, where this curve reflects the difference between environmental temperature and the temperature requirements for plant development. Most commonly, this is quantified using growing degree days (GDD) which represents the accumulation of thermal time using daily temperatures. The specific implementation varies across studies however, thus the DAESIM2-Plant model provides options for modeling thermal time.

In the simplest and most widely used form there is a linear increase in thermal time above a minimum threshold and up to a maximum threshold limit. However, the implementation and therefore interpretation differs (McMaster and Wilhelm, 1997). The first linear method compares the daily average temperature, $T_{avg}$, the base temperature and then calculates daily thermal time, $DTT$, as:

$$DTT = \begin{cases} 0 & \text{if } T_{avg} < T_b \\ T_{avg} - T_b & \text{if } T_b \leq T_{avg} \leq T_u \\ T_u - T_b & \text{if } T_{avg} > T_u \end{cases} \tag{1}$$

Where $T_{avg}$ is assumed to equal the average of the daily minimum temperature, $T_{min}$, and daily maximum temperatures, $T_{min}$, $T_b$ is the minimum threshold temperature or "base" temperature (degrees Celsius) below which the accumulation of heat units is zero, $T_u$ is the maximum threshold temperature or "upper" temperature (degrees Celsius) above which $DTT$ is kept constant. This follows the "Method 1" with an upper threshold as described in McMaster and Wilhelm (1997). The second linear method compares the $T_{min}$ and $T_{max}$ to the threshold temperatures ($T_b$, $T_u$) first, makes adjustments if necessary and then calculates $DTT$:

$$DTT = \begin{cases} 0 & \text{if } T_{avg} < T_b \\ T'_{avg} - T_b & \text{if } T_b \leq T_{avg} \leq T_u \\ T_u - T_b & \text{if } T_{avg} > T_u \end{cases} \tag{2}$$

If $T_{max} < T_b$, then $T_{max} = T_b$, if $T_{max} > T_u$, then $T_{max} = T_u$, if $T_{min} < T_b$, then $T_{min} = T_b$, if $T_{min} > T_u$, then $T_{min} = T_u$. As opposed to Method 1, the adjusted average temperature, $T'_{avg}$, using in Eq. 2 is calculated after the conditional adjustments are made to $T_{min}$ and $T_{max}$. This follows the "Method 2" with an upper threshold as described in McMaster and Wilhelm (1997).

A third, more advanced method uses hourly temperature, $T_h$, and accounts for non-linearity in the accumulation of thermal time. This introduces a concept of a thermal optimum, $T_{opt}$, at which point the thermal time is at a maximum. The approach uses a beta distribution function to relate developmental rate to temperature (Yan and Hunt, 1999; Zhou and Wang, 2018). The hourly thermal time, $HTT$, is calculated as:

$$\mathrm{HTT} = \begin{cases} 0 & \text{if } T_h < T_b \\ \left(\frac{T_h - T_b}{T_{opt} - T_b}\right)\left(\frac{T_u - T_h}{T_u - T_{opt}}\right)^{\frac{T_u - T_{opt}}{T_{opt} - T_b}} & \text{if } T_b \leq T_h \leq T_u \\ 0 & \text{if } T_u < T_h \end{cases} \tag{3}$$

Thermal time will be maximum when $T_h = T_{opt}$ and can account for an asymmetrical response above and below the optimum. The $DTT$ is then calculated as the average of $HTT$ over the day:

$$DTT = \frac{\sum_{i=1}^{n} \mathrm{HTT}_i}{n} \tag{4}$$

Where $n$ is the number of time steps per day. Applying the non-linear model (Eq. 3) requires hourly temperature data. Consequently, when DAESIM2 is run on a daily time-step with only daily minimum and maximum temperatures available, we need to assume a diurnal temperature cycle. We use the "WAVE Model" from Bal et al. (2023) to calculate a synthetic diurnal temperature profile based on daily minimum and maximum temperatures. This model combines two cosine functions: one to describe daytime warming and another for nighttime cooling. Daytime warming begins at sunrise, assumed to coincide with the daily minimum temperature, and peaks at 14:00, the assumed time of maximum temperature, which is followed by nighttime cooling that continues through to sunrise on the next day.

The choice of method to simulate thermal time in DAESIM2 depends on the available inputs and intended application. When using prescribed GDD requirements from the literature or field studies, it is essential that the model's thermal time method aligns with the method used to calculate these requirements to avoid significant errors (McMaster and Wilhelm, 1997). The linear methods are expected to perform best at intermediate temperatures and there tend to be well-documented GDD requirements for various crop types following these methods (e.g. Celestina et al., 2023). The non-linear method is expected to provide a more realistic representation of the underlying biological processes (Bonhomme, 2000), especially when representing a population of plants. For simulations across more extreme environmental conditions or future climate scenarios, the non-linear method may be preferable as it offers a more robust and constrained temperature response. However, this approach may require re-calibration for specific crop types.

#### 2.2 Germination and Emergence

From sowing time, the seed must germinate and achieve emergence, at which point the plant shifts into a photosynthetically active state. This phase is represented relatively simply with a GDD requirement that is modified by a sowing-depth. The sowing depth scaling factor decreases linearly with increasing sowing depth and ensures no germination occurs if the sowing depth exceeds a specified maximum. Thus, sowing seeds deeper will prolong the germination phase up to a limit, at which point emergence cannot occur:

$$f_{germ} = \begin{cases} 0 & \text{if } d_{sowing} < d_{sowing}^{max} \\ (1 - k_{sowdepth} \cdot d_{sowing}) & \text{else} \end{cases} \tag{5}$$

This function modifies the GDD accumulation rate during the phase that defines sowing to germination to emergence,. During all other developmental phases $f_{germ}$ is equal to 1.

#### 2.3 Vernalization

In certain crops cold temperatures mediate development. For example, in many wheat varieties the initiation and timing of flowering is moderated by prolonged exposure to cold temperatures. In a vernalization sensitive plant, exposure to cold temperatures (that satisfy the plants vernalization sensitivity) will reduce the thermal time required to reach certain developmental stages, in other words it will accelerate development. Conversely, if sufficient cold temperatures are not achieved then the accumulation of thermal time and development are slowed down. For example, wheat has vernalization genes (*Vrn-1*, *Vrn-2*, *Vrn-3*, *Vrn-4*) that control the vernalization initiation and response over time. This is a useful adaptation for the plant to adjust its development to the local environment. For winter cereals, vernalization plays a key role in preventing cold injury to reproductive tissues (Smith and Zhao, 2016).

We apply the non-linear thermal time model (Eq. 3) to determine vernalization response. This is effectively the same as Equations 6 and 7 in Wang and Engel (1998), also implemented in Equations 3 and 4 in Streck et al. (2003, their Equation 3 has a misprint in the formula), albeit with a slightly smoother response at the lower and upper temperature limits.

As described in Streck et al. (2003) "The duration of the exposure to vernalizing temperatures is measured as $VD$. One $VD$ is attained when the plant is exposed to the optimum temperature for vernalization for a period of 1 d (24 h). As temperature departs from the optimum, only a fraction of 1 $VD$ is accumulated by the plant at a given calendar day (Hodges and Ritchie, 1991; Ritchie, 1991)."

The effective vernalization days $VD$ can be calculated from the non-linear thermal time model for $HTT$ by multiplying the beta distribution function by the $(T_{opt} - T_b)$ which normalizes the curve to range between 0-1 as follows:

$$HTT = \begin{cases} 0 & \text{if } T_h < T_b \\ \left(\frac{T_h - T_b}{T_{opt} - T_b}\right)\left(\frac{T_u - T_h}{T_u - T_{opt}}\right)^{\frac{T_u - T_{opt}}{T_{opt} - T_b}} (T_{opt} - T_b) & \text{if } T_b \leq T_h \leq T_u \\ 0 & \text{if } T_u < T_h \end{cases} \tag{6}$$

Then, $HTT$ is converted to $DTT$ using Eq. 4 and this is equated to $VD$. Each day, the change in effective vernalization days is computed as follows:

$$\Delta VD = DTT \tag{7}$$

The vernalization state of the plant is determined by the accumulated effective vernalization days since the start of the current developmental phase.

The vernalization state is used to determine the vernalization factor $f_V$, which can be calculated assuming a linear or non-linear sigmoidal response function. The standard linear response function follows that of Wang and Engel (1998):

$$f_V = min\left(1, max(0, \frac{VD - Vnb}{Vnd - Vnb})\right) \tag{8}$$

The non-linear model follows that of Streck et al. (2003):

$$f_V = \frac{VD^n}{(VD_{50})^n + (VD)^n} \tag{9}$$

Where $VD_{50}$ is the vernalization state where the vernalization fraction is 50%, and $n$ is the vernalization sigmoid function shape parameter. Default values are $n$=5 and $VD_{50}$=22.5.

As an optional alternative, we also implement the linear vernalization response model from APSIM-Wheat. This linear model assumes that the vernalization factor does not start at zero. Instead, even when the vernalization state is low, $f_V$ will not wholly limit development.

$$f_V = 1 - (0.0054545R_v + 0.0003)(50 - VD) \tag{10}$$

Where $R_v$ is a vernalization sensitivity parameter, and the factor 50 is the vernalization state at full vernalization. We generalise this so that the factor 50 is set to $2 \times VD_{50}$, which is functionally the same definition as $VD_{50}$ in the non-linear model.

#### 2.4 Developmental Rate

The impact of sowing depth and vernalization on plant development is to moderate the plant developmental rate. As such, the germination factor, $f_{germ}$, and the vernalization factor, $f_V$, are applied as modifiers on the $GDD$ state variable during specific developmental phases:

$$\frac{dGDD}{dt} = DTT \cdot f_V \cdot f_{germ} \tag{11}$$

Where $GDD$ is the growing-degree-days, which tracks thermal time in the plant, $DTT$ is the daily thermal time. When running the model, the $GDD$ state is set to zero at the time of sowing.

#### 2.5 Developmental Phase Details

On top the what has been described above, some developmental phases have specific processes associated with them.

- Grain production: Grain production occurs during a specific developmental phase, typically following flowering and preceding maturity/senescence. Grain production is handled in different ways depending on the crop type (described further below).

- Maturity: The maturity phase in annual crops typically follows reproduction. During maturity the plant undergoes age-related senescence. During this phase of the life cycle the crop undergoes transformations to source-sink dynamics, hormone signalling (see Gregersen et al., 2013). Photosynthesis is down-regulated in part by nitrogen retranslocation out of the leaves and into the reproductive organs. Hydraulic changes in the xylem and leaves can also induces progressive down-regulation of plant function, including photosynthesis (e.g. Locke and Ort, 2014). There is generally a coordinated reduction in photosynthesis and transpiration during maturity. This is represented in DAESIM2-Plant by progressively down-regulating key process parameters related to photosynthetic capacity ($V_{cmax}^{25}$), stomatal conductance ($g_1$) and plant hydraulic conductance ($k_{rl}$). Mathematically, they are linearly scaled down from their value at the beginning of maturity to zero by the end of maturity. This affects the source supply of assimilates. As this occurs, both autotrophic respiration and turnover (e.g. leaf drop) continue at their nominal rates.

### 3 Leaf Gas Exchange

The leaf gas exchange model simulates the exchange of carbon dioxide and water vapor at the leaf surface. It simulates C3 photosynthesis using the mechanistic model of Farquhar et al. (1980) where key processes are related to environmental drivers including absorbed light, temperature and the concentration of atmospheric CO2 (Farquhar et al., 1980). Stomatal conductance is modeled using the theory of optimal stomatal behaviour, whereby stomata act to maximize carbon gain through photosynthesis while minimizing water loss through transpiration (Cowan and Farquhar, 1977). When coupled with the mechanistic model for photosynthesis, this model reliably describes coupled photosynthesis-stomatal conductance behaviour across a range of environmental conditions (Medlyn et al., 2011).

The model includes key biochemical parameters such as the maximum carboxylation rate ($V_{cmax}$), the maximum electron transport rate ($J_{max}$), and mesophyll and stomatal conductance parameters. It accounts for temperature-dependent kinetics, light-driven electron transport, and the balance between Rubisco-limited, electron transport-limited, and triose phosphate utilization-limited photosynthesis.

#### 3.1 C3 Photosynthesis

The mechanistic basis of the C3 photosynthesis model has been described in detail elsewhere (Farquhar et al., 1980; von Caemmerer, 2000; Yin et al., 2021). It has been adopted widely for interpreting leaf gas exchange measurements, describing the controls and limits of photosynthesis, and predicting ecophysiological processes from canopy to global scales as a component of Earth System Models. Here, we show the fundamental equations, some minor modifications to the original Farquhar et al. (1980) model and the implementation in DAESIM2.

The original model predicts gross photosynthetic rate, $A_g$, as the minimum of two potentially limiting processes, the Rubisco-limited rate and the electron transport-limited rate. A third process was included in the model later which represents the triose phosphate utilization-limited rate, which is most important under high CO2 concentrations. Net photosynthesis is expressed as $A_g$ minus the day respiration rate, $R_d$, which represents the mitochondrial CO2 release not including photorespiration:

$$A_n = min\{A_c, A_j, A_p\} - R_d = A_g - R_d \tag{12}$$

The Rubisco-limited rate is expressed as:

$$A_c = V_{cmax} \cdot \frac{C_i - \Gamma^*}{C_i + K_C \cdot (1 + \frac{O_i}{K_O})} \tag{13}$$

Where $V_{cmax}$ is the maximum carboxylation rate, $C_i$ is the CO2 partial pressure in the intercellular spaces, $\Gamma^*$ is the CO2 compensation point in the absence of day respiration, $K_C$ and $K_O$ are the Michaelis-Menten constants for CO2 and O2, respectively, and $O_i$ is the partial pressure of O2 in the intercellular spaces. The $\Gamma^*$ is calculated as:

$$\Gamma^* = \frac{1}{2} \cdot \frac{O_i}{S} \tag{14}$$

Where $S$ is the relative CO2/O2 specificity factor for Rubisco. In theory, $S$ is equal to $V_{cmax}/V_{omax} \cdot K_O/K_C$ but it is often used as a separate parameter.

The electron transport-limited rate is expressed as:

$$A_j = \frac{J}{4} \cdot \frac{C_i - \Gamma^*}{C_i + 2\Gamma^*} \tag{15}$$

Where $J$ is the electron transport rate, calculated using a non-rectangular hyperbola model as a function of absorbed irradiance:

$$J = \frac{\alpha Q_{abs} + J_{max} - \sqrt{(\alpha Q_{abs} + J_{max})^2 - 4\theta\alpha Q_{abs} J_{max}}}{2\theta} \tag{16}$$

Where $J_{max}$ is the maximum electron transport rate, $Q_{abs}$ is the absorbed irradiance, $\alpha$ is the quantum yield of electron transport and $\theta$ is an empirical curvature parameter.

The triose phosphate utilization-limited rate is expressed as:

$$A_p = 3 \cdot TPU \cdot \frac{C_i - \Gamma^*}{C_i - (1 + 3\alpha_g)\Gamma^*} \tag{17}$$

Where $TPU$ is the triose phosphate utilization rate and $\alpha_g$ is the fraction of glycolate not returned to the chloroplast (Eq. 7 in Ellsworth et al., 2015).

The $R_d$ is commonly assumed to scale with $V_{cmax}$ given their shared dependence on leaf nitrogen (Atkin et al., 2015). Thus, $R_d$ at 25 degrees Celcius is represented as a fraction of $V_{cmax}$ at 25 degrees Celsius as follows:

$$R_d^{25} = R_{ds} \cdot V_{cmax} \tag{18}$$

Where $R_{ds}$ is the scaling coefficient.

#### 3.2 Temperature Dependencies

Many of the kinetic parameters of the C3 photosynthesis model are temperature sensitive. There are three temperature response functions (see Medlyn et al., 2002), which apply to different parameters depending on their assumed temperature response. The Arrhenius temperature function is given by:

$$k(T) = k_{opt} \cdot exp^{((E_a(T - T_{opt}))/(T_{opt} \cdot R \cdot T))} \tag{19}$$

Where $k_{opt}$ is the parameter value at the optimum temperature (typically 25°C), $T$ is the leaf temperature (K), $E_a$ is the activation energy, $R$ is the universal gas constant and $k(T)$ is the temperature-adjusted parameter. The peaked-Arrhenius temperature response function is given by:

$$k(T) = k_{25} \cdot exp^{\left(\frac{E_a(T-298.15)}{(298.15 \cdot R \cdot T)}\right)} \cdot \frac{(1 + exp^{(298.15 \cdot \Delta S - H_d)/(289.15 \cdot R)})}{(1 + exp^{(T \cdot \Delta S - H_d)/(T \cdot R)})} \tag{20}$$

Where $k_{25}$ is the parameter at 25°C, $H_d$ is the deactivation energy and $\Delta S$ is the entropy term. The $Q_{10}$ temperature function is given by:

$$k(T) = k_{ref} \cdot Q_{10}^{\left(\frac{T - T_{ref}}{10}\right)} \tag{21}$$

Where $k_{25}$ is the parameter at the reference temperature, $T_{ref}$, and $Q_{10}$ is the temperature sensitivity coefficient.

#### 3.3 Stomatal Conductance

The representation of stomatal conductance follows a unified approach that describes an empirical expression which is consistent with optimal stomatal behaviour, as first described by Cowan and Farquhar (1977). Optimal stomatal behaviour theory postulates that stomata should act to maximize carbon gain (photosynthesis, $A$) while at the same time minimizing water lost ($E$, transpiration). The model follows that described in Medlyn et al. (2011) and Medlyn et al. (2012) as implemented in (Duursma, 2015).

The model (Eq. 11 in Medlyn et al., 2011), note the corrigendum in Medlyn et al. (Eg. 1 in 2012), expresses the stomatal conductance of water vapour, $g_{sw}$, as:

$$g_{sw} \approx g_0 + 1.6(1 + \frac{g_1}{\sqrt{D}})\frac{A_n}{C_a} \tag{22}$$

when $D$ is the leaf-to-air vapor pressure deficit (kPa), $C_a$ is the atmospheric CO2 partial pressure, $g_0$ and $g_1$ are fitting parameters, where $g_1$ has units of kPa$^{0.5}$, which can be shown to be proportional to the marginal water cost of carbon gain (often denoted as $\lambda$ in the optimal stomatal theory) and with the CO2 compensation point, $\Gamma*$ (Medlyn et al., 2011). Here, we include a term to describe the down-regulation of stomatal conductance in relation to plant water status, specifically leaf water potential, using the so-called "Tuzet" model (Tuzet et al., 2003). The Tuzet model describes the stomatal response to leaf water potential with a logistic-type function as follows:

$$f_{sv} = \frac{1 + exp^{(s_f \cdot \Psi_f)}}{1 + exp^{(sf(\Psi_f - \Psi_l))}} \tag{23}$$

Where $\Psi_L$ is the bulk leaf water potential (MPa), $s_f$ is the stomatal sensitivity factor ($\text{MPa}^{-1}$) and $\Psi_f$ is a reference leaf water potential at which point there is 50% down-regulation (i.e. $f_{sv}$=0.5). The factor $f_{sv}$ ranges between 1 (no stomatal down-regulation) and 0 (complete stomatal down-regulation) as $\Psi_L$ declines. Incorporating the Tuzet model into the stomatal conductance expression gives:

$$g_{sw} = g_0 + 1.6(1 + f_{\Psi_l} \cdot \frac{g_1}{\sqrt{D}})\frac{A_n}{C_a} \tag{24}$$

Certain cultivars of wheat are known to exhibit different stomatal responses to leaf water potential, due to differences in xylem vulnerability to cavitation under water stress. For example, cv Armstrong exhibits stomatal closure at lower xylem pressures than cv Cesario (Fig. 3b in Torres-Ruiz et al., 2024). Also in wheat, measurements show that there is about 50% stomatal closure at a water potential of -0.85 MPa and full stomatal closure by about -2.0 MPa (Corso et al., 2020), providing a basis to parameterize the Tuzet model. If accounting for whole-plant hydraulic limitations and xylem cavitation in this model, it may be more suitable to set a $\Psi_f$ lower, between -2.4 MPa and -2.2 MPa (Corso et al., 2020).

#### 3.4 Coupling Photosynthesis and Stomatal Conductance

The equations above define the relationship between the net assimilation rate, $A_n$, and the intercellular CO2 concentration ($C_i$), and the relationship between the stomatal conductance, $g_{sw}$, and net assimilation rate. A third equation defines the diffusional supply of CO2 for photosynthesis following Fick's law, $A_n$ to $g_{sw}$ via the CO2 concentration gradient between the inside and outside of the leaf:

$$A_n = \frac{g_{sw}}{1.6} \cdot (C_i - C_a) \tag{25}$$

Together, these form a system of three equations with three unknowns: $A_n$, $C_i$, and $g_{sw}$. At steady state, CO2 supply via Fick's law (Eq. 25) and demand due to photosynthetic metabolism must be balanced, with equilibrium at the intersection of supply and demand curves as a function of $C_i$. To solve this system, Arneth et al. (2002) derived two quadratic expressions for optimal $C_i$, corresponding to the Rubisco-limited and electron transport-limited rates. Medlyn et al. (2011) further extended this framework within the unified optimal stomatal conductance model, which we implement following Duursma (2015), allowing $C_i$ to be solved through these quadratic formulations.

### 4 Plant Hydraulics

The plant hydraulics model represents the hydraulic pathway extending from the soil, through the plant, and to the transpiring leaves. Accumulating empirical evidence supports a strong association between stomatal conductance, leaf water status, and the hydraulic properties of the plant–soil system (Bonan et al., 2014; Mencuccini, 2003; Choat et al., 2012; Manzoni et al., 2013), including in agricultural species like wheat (Ranawana et al., 2021). In DAESIM2-Plant, this is conceptualized as a soil-plant-atmosphere continuum that adheres to Darcy's Law for steady-state water flow driven by a water potential gradient. This gradient forms between the soil and canopy as water moves upward, supplied by the roots and is ultimately lost through transpiration. Gravitational potential is neglected as it only significantly impacts conductance over long pathways, such as in tall trees. Additionally, the model excludes the effect of osmotic pressure from dissolved solutes. The hydraulic pathway is treated as one-dimensional and steady-state, simplifying the representation of water movement through the plant and ignoring dynamics of water storage capacity.

The soil water potential, $\Psi_s$ (MPa), is determined from the volumetric soil water content based on a power function (Campbell, 1974):

$$\Psi_s = \Psi_e \cdot \left(\frac{\theta}{\theta_{max}}\right)^{-b_{soil}} \tag{26}$$

Where $\theta$ is the volumetric soil water content (m3 m-3), $\theta_{max}$ is the volumetric soil water content (m3 m-3) at saturation, $\Psi_e$ is the saturating value of soil water potential (MPa), and $b_{soil}$ is an empirical soil-water retention curve parameter (soil-type specific). This function is constrained to values where $\theta \leq \theta_{max}$. The parameters can be estimated from a soil moisture retention curve (Campbell, 1974) and are related to the physical properties of the soil (Cosby et al., 1984).

The soil hydraulic conductivity, $K_s$ (mol m-1 s-1 MPa-1; mol H2O per m per second per MPa pressure difference) is dependent on $\Psi_s$ as follows:

$$K_s = K_{sat} \cdot \left(\frac{\Psi_e}{\Psi_s}\right)^{(2+3/b_{soil})} \tag{27}$$

Where $K_{sat}$ is the saturating value of soil hydraulic conductance (mol m-1 s-1 MPa-1; mol H2O per m per second per MPa pressure difference).

The soil-to-root radial hydraulic conductivity is represented by:

$$K_{sr} = K_s \cdot \frac{L_v}{k_{sr,coeff}} \tag{28}$$

Where $L_v$ (g d.wt root m-3 soil) is density of conducting roots, calculated as:

$$L_v = \frac{W_R \cdot f_r}{d_{soil}} \tag{29}$$

Where $W_R$ is the total root biomass (g d.wt m-2), $d_{soil}$ is the soil layer depth (m) and $f_r$ is the fraction of $W_R$ in the given soil layer. The parameter $k_{sr,coeff}$ is an empirical constant that describes the relationship between root biomass and the conducting surface area of the roots (g d.wt-1 m-1), which is represented differently across the literature. In principle, root conducting capacity is influenced by properties such as root hairs, tapering, clustering, root radius, root length (in combination these govern bulk properties such as root length index and/or root area index), and the distance from the root surface to bulk soil water, as well as biophysical changes like root shrinkage during drying. Commonly, the "single root cylinder" model is used, which conceptualizes roots as vertical cylinders with a fixed radius drawing water from a surrounding soil cylinder (Gardner, 1960), which has been implemented in many soil-plant-atmosphere continuum models (Williams et al., 2001; Katul et al., 2003; Duursma et al., 2008; Duursma and Medlyn, 2012; Bonan et al., 2014). Nobel (2009) provides a fundamental description of this model. Although useful, the single root model assumes homogeneity in root properties, orientation, and distribution, and requires that properties like root radius and length index be specified independently, which is often impractical without highly detailed field data. In DAESIM2-Plant, instead of defining fixed values for these parameters, we simplify them into a single empirical parameter that encapsulates their combined properties. This approach reflects the behavior described by the single root cylinder model, as $k_{sr,coeff}$ can be easily derived from it. Justification for this simplification comes from modeling studies indicating that water uptake functions based on root dry mass, length, or surface area density yield similar results (Himmelbauer et al., 2008).

The soil-to-root radial hydraulic conductivity, $K_{sr}$, is converted to a leaf-area specific soil-to-root hydraulic conductance (mol m-2 s-1 MPa-1) as follows:

$$k_{sr,l} = K_{sr}/LAI \tag{30}$$

Where $LAI$ is the canopy leaf area index (m2 one-side leaf m-2 ground).

The total leaf-area specific conductance from the soil to the leaf, $k_{tot,l}$ (mol m-2 s-1 MPa-1), including root-to-leaf conductance, $k_{rl,l}$ (mol m-2 s-1 MPa-1), is calculated assuming a one-dimensional pathway in series that follows Ohm's Law for the hydraulic conductances as follows:

$$\frac{1}{k_{tot,l}} = \frac{1}{k_{sr,l}} + \frac{1}{k_{rl,l}} \tag{31}$$

Where $k_{rl,l}$ is assumed constant (see Bonan et al., 2014). This parameter has been be measured in some crop types (Tsuda and Tyree, 2000).

#### 4.1 Balancing Plant Transpiration and Root Water Uptake

It is assumed that the rate of water loss from the canopy via transpiration is balanced by the rate of plant water uptake from the soil (Meinzer, 2002; Bonan et al., 2014), thus, the plant water balance is in steady-state. Given this, the transpiration rate can be described in terms of plant water uptake through the soil-to-leaf hydraulic continuum. Following Darcy's law, the leaf-area specific transpiration rate, $E_l$, is given by the hydraulic conductance along the soil-to-leaf pathway multiplied by the water potential gradient:

$$E_l = k_{sr,l} \cdot (\Psi_s - \Psi_r) = k_{rl,l} \cdot (\Psi_s - \Psi_r) = k_{tot,l} \cdot (\Psi_s - \Psi_l) \tag{32}$$

Alternatively, the leaf-area specific transpiration rate, $E_l$, can be described using the total diffusional resistance between the inside of the leaf and the surrounding air. This resistance includes both the stomatal resistance to water vapour ($r_{sw}$) and the leaf boundary layer resistance to water vapour ($r_{bw}$), as defined by Fick's law:

$$E_l = \frac{(W_i^w - W_a^w)}{r_{sw} + r_{bw}} = \frac{VPD}{r_{sw} + r_{bw}} \tag{33}$$

Where both $r_{sw}$ and $r_{bw}$ are defined below. In this equation, $W_i^w$ represents the water vapor partial pressure inside the leaf, and $W_a^w$ is the water vapor partial pressure outside the leaf, both expressed in molar concentration (mol m-3). Since the leaf's intercellular spaces are generally assumed to be saturated with water vapor, $W_i^w$ is equivalent to the saturation vapor pressure, allowing $VPD$ (vapor pressure deficit, in molar concentration) to be used directly in this expression. Note that in the coupled leaf photosynthesis-stomatal conductance model, the leaf boundary layer resistance is ignored, but it is included in transpiration rate. This may require updating in future but would require a re-write of the quadratic expressions to solve for $C_i$ (e.g. Bonan et al., 2014).

The stomatal resistance to water vapour, $r_{sw}$ (m s-1), is proportional $g_{sw}$ as follows:

$$r_{sw} = \frac{1}{g_{sw}} \cdot \frac{P}{R_w \cdot T} \tag{34}$$

Where $P$ is the air pressure (Pa), $R_w$ is the specific gas constant for water vapor (J mol-1 K-1), and $T$ is the temperature (K). The combined term on the right-hand-side makes the appropriate unit conversions from $g_{sw}$ to $r_{sw}$.

The leaf boundary layer resistance to water vapor, $r_{bw}$, depends on the thickness of the boundary layer, $\delta_{bl}$, and the binary diffusion coefficient of water vapor in air, $D_{H2O}$ (approximately 2.42 x 10-5 m2 s-1 at 20 degrees C). Experiments show that $\delta_{bl}$ depends on both leaf size and wind speed, following a power law relationship:

$$r_{bw} = \frac{\delta_{bl}}{D_{H2O}} = k_{wl} \cdot \sqrt{\left(\frac{d_{leaf}}{u}\right)} \cdot \frac{1}{D_{H2O}} \tag{35}$$

Where $u$ is the wind speed (m s-1), $d_{leaf}$ (m) is the leaf dimension parameter which represents the average length of the leaf in the downwind direction-and the wind speed, and $k_{wl}$ ($s^{1/2}$ m-1) is an empirical coefficient, which typically ranges from 0.004 to 0.006 (Nobel, 2009). This relationship is assumed valid for wind speeds exceeding 0.05 m s-1. For wind speeds below this threshold, $r_{bw}$ is treated as a constant (at $u = 0.05$ m s-1) to prevent it from approaching infinity as wind speed nears zero.

With Eqs. 32 and 33, we have two expressions for $E_l$, both of which are functions of the bulk leaf water potential, $\Psi_l$, noting that Eq. 33 includes $\Psi_l$ in its expression for $r_{sw}$ (Eq. 34) and the coupled photosynthesis-stomatal conductance model that determines $g_{sw}$ (Eq. 24). Given that we assume the plant water uptake and plant water loss are in steady-state, we use a numerical bisection method that determines the value of $\Psi_l$ that balances these two equations to within an acceptable error tolerance. Given $\Psi_l$, the actual canopy photosynthesis and transpiration rate can be calculated with Eqs. X and Y, respectively.

#### 4.2 Root Depth and Vertical Distribution

Root depth and root vertical distribution are crucial plant traits that determine a plant's ability to access soil resources, such as water and nutrients. These traits influence water uptake, especially under varying soil moisture conditions, directly affecting plant function and overall resilience to environmental stress.

To represent this in DAESIM2-Plant, we assume root density has a defined vertical distribution from the surface down to a specified rooting depth.

The plant rooting depth evolves dynamically over the growing season, constrained by a maximum potential rooting depth. Rooting depth, $d_r$ (m), is determined as the minimum of the two potential rooting depths, one based on developmental rate, $d_r^{dev}$ (m), and the other based on root biomass, $d_r^W$ (m), as follows:

$$d_r = min\{d_r^{dev}, d_r^W\} \tag{36}$$

The potential rooting depth based on development, $d_r^{dev}$ (m), is assumed to be a linear function of the developmental rate from germination up to a defined point in the growing season where the maximum can be achieved. This is sometimes quantified with the so-called root penetration rate (Kirkegaard and Lilley, 2007). The model calculates this by:

$$d_r^{dev} = d_{r,max} \cdot \frac{GDD(t)}{GDD_{d_r}^{max}} \tag{37}$$

Where $d_{r,max}$ (m) is the maximum potential rooting depth, $GDD(t)$ is the current growing-degree day state of the plant and $GDD_{d_r}^{max}$ is the required $GDD$ that the plant can reach the maximum potential rooting depth, typically assumed to be the beginning of anthesis in most crops. The potential rooting depth based on root biomass is calculated by:

$$d_r^W = W_R \cdot SRD \tag{38}$$

Where $W_R$ (g d.wt m-2) is the total root biomass and $SRD$ (m g d.wt-1) is the specific root depth which represents the average ratio of root depth to total root dry biomass.

Multiple studies have demonstrated that the distribution of roots in the soil profile, measured as root mass density (g root m-3 soil) or root length density (m root m-3 soil), declines exponentially or near-linearly with depth from the soil surface (see Fan et al., 2016; Haberle and Svoboda, 2014, and references therein). Also see Osborne et al. (2020). Schenk and Jackson (2002) used a logistic dose–response curve to realistically represent the cumulative root distribution which was adapted by Fan et al. (2016) and further evaluated to describe root distributions in common temperate crops. This equation is given by:

$$\frac{R}{R_{max}} = \frac{1}{1 + (d/d_{50})^c} \tag{39}$$

where $R$ is the cumulative amount (i.e., biomass mass or root length) of roots to soil depth $d$ (m; i.e. the amount of roots above profile depth $d$), $R_{max}$ is the total amount of roots, $d_{50}$ is depth at which 50% of total root amount has accumulated and $c$ is a dimensionless shape parameter. The above equation is not constrained to rooting depth, as it continues to be positive definite to infinite soil depth. This can be modified so that all roots are confined to the actual rooting depth by doing the following:

$$\frac{R}{R_{max}} = Y(d) = \frac{1}{1 + (d/d_{50})^c} + \left(1 - \frac{1}{1 + (d_r/d_{50})^c}\right) \cdot \frac{d}{d_r} \tag{40}$$

Where $Y(d)$ is the cumulative fraction of roots from the soil surface to depth $d$ (m). This function is constrained to values where $d \leq d_r$. Fan et al. (2016) provide values for the parameters above for eleven different crop types, including wheat and canola. This provides a continuous vertical profile for root biomass as a function of soil depth.

***Parameterization notes:

For maximum rooting depth across crop species see Merrill et al. (2002). For determining the developmental phase (in growing degree days) where roots achieve their maximum rooting depth, see Smit and Groenwold (2005).

#### 4.3 Implementation in a Multi-Layer Soil

Soil moisture availability and root density both vary with depth in the soil profile which are important factors in determining the plants access to soil water. In DAESIM2-Plant, the soil is divided into $n$ discrete layers, each with user-defined depth and properties that affect soil-to-root hydraulics. Soil moisture, including its vertical distribution, is prescribed in the DAESIM2-Plant model and is not further discussed here. For the vertical distribution of roots, the user-defined discretization of the soil profile determines the depth to the bottom of each layer, which allows for the calculation of the cumulative fraction of roots from the surface to that layer with Eq. 40. The fraction of roots, $f_r$, in soil layer $n$ is then calculated as the difference between $Y$ at the bottom of layer $n$ and $Y$ at the bottom of layer $n-1$, with the exception of the uppermost soil layer which is simply the first value of $Y$ i.e. $Y(n=0)$. The calculated $f_r$ per layer is used in Eq. 29 to determine the root biomass per soil layer.

The soil-to-root hydraulic conductivity, $K_{sr}$, is calculated for each soil layer to capture vertical variation in soil moisture availability and root biomass. Similar to determining a bulk canopy leaf water potential, we calculate a bulk root-zone $K_{sr}$ and a bulk root-zone soil water potential, $\Psi_s$, which are used as the inputs for the root water uptake and transpiration rate equations (see Eq. 32). To calculate these bulk values, we assign a weight to each soil layer, proportional to the layer's soil-to-root hydraulic conductivity $K_{sr}$. Since $K_{sr}$ depends on both root biomass and moisture availability in each layer (see Eq. 28), this weighting approach inherently accounts for these factors. For instance, layers without root biomass receive a weight of zero, or layers with higher soil moisture receive relatively larger weighting. The bulk root-zone soil water potential $\Psi_s$ and soil-to-root hydraulic conductivity $K_{sr}$ are then computed as a weighted average, where each layer-specific value is multiplied by its respective weight and then the sum is taken over all layers.

### 5 Canopy Radiative Transfer

The scattering and absorption of radiation in the canopy is described using the two-stream approximation of radiative transfer theory. Two-stream radiative transfer theory applies fundamental radiative transfer principles to represent light interactions within a turbid medium, which in the case of vegetation canopies includes leaves, stems and/or the soil surface. Other approaches to modeling canopy radiative transfer, such as a big-leaf model or a sunlit-shaded two big-leaf model, are computationally efficient but make a number of assumptions about the vertical distribution of light, canopy optical properties and canopy physiological properties that are not always supported by data (Bonan et al., 2021). These simpler schemes often lead to biases in modeled carbon, water or energy fluxes (Luo et al., 2018; Wang and Frankenberg, 2022).

Fundamentally, the canopy radiative transfer model is based on the radiative transfer equation, which describes the change in radiance as radiation is absorbed, emitted, and scattered through a medium. The two-stream approach simplifies the full equation by reducing the directional complexity of radiative fluxes to two primary streams: a backward scattering hemisphere (reflected) and a forward scattering hemisphere (transmitted). The approximation assumes that the incoming sky diffuse radiation and the scattered radiation within the canopy are isotropic in inclination, while direct beam maintains its directional properties. The approach ensures energy conservation, distributing absorbed, transmitted, and reflected radiation through the canopy and down to the soil surface. This simplification leverages assumptions about scattering symmetry and within-layer homogeneity to make the model computationally feasible while maintaining its physical accuracy. Originally implemented to represent the land-surface and vegetation canopies as a single turbid layer (Dickinson, 1983; Sellers, 1985) this approach has been adopted widely in representing vegetation canopies and in the land surface component of climate models.

We use the two-stream approximation as implemented in the multilayer canopy model by Bonan et al. (2018) and extended by Bonan et al. (2021) to handle any number of layers, as in the Community Land Model multi-layer version 1 (CLM-ml v1). Each canopy layer represents a turbid medium with horizontally homogeneous properties. The properties of each layer are characterized by structural properties, such as leaf area index (LAI), stem area index (SAI), leaf angle distribution, and foliage clumping, alongside optical properties like transmittance and reflectance, along with soil albedo at the base. This is useful as vertical variation in canopy structure and optical properties is represented explicitly, which can be important for many canopies including crops which have different vertical structures across species and during different developmental stages (Hosoi et al., 2009; Li et al., 2015). At each layer, incident radiation is partitioned into direct beam and diffuse components, both handled separately to capture differential effects within the canopy. Direct beam radiation illuminates sunlit foliage due to its directional nature, while diffuse radiation is distributed more evenly and illuminates both sunlit and shaded foliage. This separation is critical for accurately estimating light absorption profiles, as each component interacts differently with the canopy structure. The sunlit and shaded portions of the canopy are determined dynamically based on canopy structure and sun position. The model calculates upward and downward radiative fluxes for sunlit and shaded portions of each layer, iteratively moving through the canopy layers and accounting for absorption and scattering at each step. This approach is essential to realistically representing vertically structured canopies in response to both direct and diffuse radiation, and provides realistic predictions of light availability, photosynthesis, and energy balance within canopies.

### 6 Plant Carbon Balance, Respiration and Allocation

#### 6.1 Plant Carbon Balance

The plant carbon balance is a function of the new assimilates from photosynthesis and losses due to respiratory processes and senescence i.e. litter production. The rate of change in plant carbon is given by:

$$\frac{d_{C_{plant}}}{dt} = GPP - R_a - S = NPP - S \tag{41}$$

Where $GPP$ is the gross primary productivity (g C m-2 d-1), $R_a$ is the total autotrophic respiration rate (g C m-2 d-1) and $S$ is the plant litter production rate (senescence; g C m-2 d-1). The net primary productivity, $NPP$ (g C m-2 d-1), is the difference between $GPP$ and $R_a$.

#### 6.2 Autotrophic Respiration

Autotrophic respiration is simulated following the growth-and-maintenance respiration paradigm (McCree, 1970), which stipulates that all respiratory metabolic processes fall under either the growth or maintenance terms (Amthor, 2000). Growth respiration is associated with the growth rate of structural biomass and maintenance respiration is temperature sensitive and associated with the mass of live biomass.

Leaf maintenance respiration, $R_{a,leaf}$, is assumed to equal the sum of the leaf mitochondrial respiration rate, $R_d$ (Eq. 18 including temperature corrections with Eq. 21) over the canopy. Root maintenance respiration is proportional to the root carbon pool size as follows:

$$R_{a,root} = f(T) \cdot m_{r,root} \cdot C_{root} \tag{42}$$

Where $C_{root}$ is the total root carbon pool (g C m-2), $m_{r,root}$ is the specific root maintenance respiration rate (d-1) and $f(T)$ is the temperature response function represented by a $Q_{10}$ function (Eq. 21). We note that maintenance respiration of stem and grain tissue is currently ignored, as these will be relatively minor terms for crops. The total maintenance respiration rate, $R_m$, is the sum of $R_{a,leaf}$ and $R_{a,root}$.

Growth respiration is represented as a fixed fraction of available assimilates after maintenance respiration has been accounted for (i.e. $GPP - R_m$), thereby assuming that maintenance of existing living tissue is prioritized by the plant before new growth. This is represented as:

$$R_g = \alpha_{R_g}(GPP - R_m) \tag{43}$$

Where $\alpha_{R_g}$ represents the proportion of available assimilates that are respired during growth, usually in the range 0.1-0.3.

#### 6.3 Optimal Trajectory Carbon Allocation

Background: There is evidence that plants modify their allocation in response to limiting factors. Poorter and Nagel (2000) conducted an extensive literature review and found that functional equilibrium theory provides a good foundation to model this adjustment of allocation in response to limiting resources and noted that the "responses to light, nutrients and water agreed with the (qualitative) prediction of the 'functional equilibrium' theory".

Also see Kulbaba et al. (2023), who discuss in detail the concept of "chasing the fitness optimum: temporal variation in the genetic and environmental expression of life-history traits for a perennial plant". They state nicely in their abstract "The ability of plants to track shifting fitness optima is crucial within the context of global change, where increasing environmental extremes may have dramatic consequences for life history, fitness, and ultimately population persistence. However, tracking changing conditions relies on the relationship between genetic and environmental variance, where selection may favour plasticity, the evolution of genetic differences, or both depending on the spatial and temporal scale of environmental heterogeneity."

The approach to carbon allocation adopted here in DAESIM2-Plant is modified from Potkay et al. (2021).

The instantaneous allocation fraction to pool $k$ is proportional to the ratio of the marginal gain divided by the marginal cost:

$$u_k \propto \frac{marginal\ gain_k}{marginal\ cost_k} \tag{44}$$

where $u_k$ is the instantaneous allocation fraction to pool $k$. The marginal gain per pool is equal to

$$marginal\ gain_k = \frac{d}{dC_k}[LAI \cdot (A_n + R_d) - R_m] = \frac{d}{dC_k}[GPP - R_m] \tag{45}$$

where $LAI$ is the leaf area index, $A_n$ is net photosynthetic rate, $R_d$ is leaf mitochondrial respiration rate, and $R_m$ is the maintenance respiration rate.

The marginal cost takes into account the mean residence time of the pool. As discussed in Potkay et al. (2021) this means that carbon allocation "considers how long any investment of carbon will last and potentially benefit a tree [plant]. Investments with short-lived payoffs (i.e. small $\tau_i$) benefit the tree [plant] only briefly and thus reflect poor investments over the duration of a tree's [plant's] life." To account for this, one may consider the instantaneous senescence rate of a given pool ($S_i$; g C m-2 d-1):

$$S_k = \frac{C_k}{\tau_k} = k_C^k\ C_k \tag{46}$$

where $C_k$ is the pool size (g C m-2) and $\tau_k$ is the mean life span of the pool (days), and $k_C^k$ is the carbon turnover rate (days-1), all of which are expressed for the respective pools ($k$). The marginal cost is calculated as the change in senescence rate divided by the change in pool size.

$$marginal\ cost_k = \frac{dS_k}{dC_k} = \frac{1}{\tau_k} = k_C^k \tag{47}$$

We note here that Potkay et al. (2021) seems to have written out the above equation incorrectly in their supplementary material (Equation S.8.3), incorrectly writing $dS_k/dC_k = tau_k$.

In practice, when the economic gain is negative, the allocation is set to zero. Furthermore, to ensure allocation fractions to all pools sum to unity, each marginal gain-cost ratio is normalized to the sum of marginal gain-cost ratios for all pools:

$$u_k = \frac{max(0, \frac{marginalgain_k}{marginalcost_k})}{\sum_j max(0, \frac{marginalgain_j}{marginalcost_j})} \tag{48}$$

where $j$ is the vector of $k$ carbon pools.

##### 6.3.1 Accounting for fixed allocation fractions to non-optimal pools

We can modify this to include constant allocation fractions for some pools. For example, we can assume that the carbon allocation to leaves and roots follows the optimal trajectory principle outlined above, but that allocation to stems ($u_S$) and grains ($u_G$) is fixed. To account for this we can do the following. First, we note that the sum of all four allocation fractions must sum to unity:

$$u_L + u_R + u_S + u_G = 1 \tag{49}$$

The allocation fractions to $u_L$ and $u_R$ are calculated using the optimal trajectory equations further above, which we denote $u'_L$ and $u'_R$, respectively, remembering that $u'_L + u'_R = 1$. To account for constant, non-zero terms for $u_S$ or $u_G$, we must scale $u'_L$ and $u'_R$ by a factor $\alpha$:

$$u_L = \alpha u'_L \tag{50}$$

and

$$u_R = \alpha u'_R \tag{51}$$

Therefore, we now have:

$$\alpha u'_L + \alpha u'_R + u_S + u_G = \alpha(u'_L + u'_R) + u_S + u_G = 1 \tag{52}$$

We know that $u'_L + u'_R = 1$, therefore:

$$\alpha(1) + u_S + u_G = 1 \tag{53}$$

and

$$\alpha = 1 - (u_S + u_G) \tag{54}$$

So, the actual allocation fractions to the leaf and root pools are:

$$u_L = (1 - (u_S + u_G))u'_L \tag{55}$$

and

$$u_R = (1 - (u_S + u_G))u'_R \tag{56}$$

This scales the optimal trajectory coefficients equally and in a way that maintains the total sum of allocation coefficients equal to 1.

### 7 Grain Production

#### 7.1 Wheat (Triticum)

The grain yield, $GY$, in wheat is determined by the number of grains generated per unit area, $GN$, and the mean individual weight of those grains, $GW$. As reviewed by Pretini et al. (2021), $GN$ is the most critical factor "as sink limitation remains present during grain filling" (Borrás et al., 2004; Acreche and Slafer, 2006; González et al., 2014). The $GN$ is established during the "critical period", which occurs approximately 20 days before and 10 days after anthesis, which corresponds closely with formation of spikes and determines the spike dry weight (at anthesis), fertile floret number and $GN$ (Fischer et al., 2024). As reviewed by Fischer et al. (2024), the critical period is defined as "the interval between flag leaf emergence and first anthesis, each across 50% of the culms in any crop, a period encompassing most of the accumulation of spike dry matter, in turn determining floret survival and final fertile floret numbers/m$^2$".

Pretini et al. (2021) stated that "after ca. 10 days post-anthesis, the number of fertile florets that set grains (grain set: GST) is established (Fischer, 1975; 1985), determining GN (graph 3 in Fig. 1A). Thus, the most validated eco-physiological model considers that the floret survival, and hence GN, depends on the partitioning of assimilates to the growing spikes (Fischer and Stockman, 1980; Kirby, 1988; Ghiglione et al., 2008; González et al., 2011a). This leads to a strong relationship between GN and spike dry weight per unit area at anthesis (SDW) usually observed (graph 4 in Fig. 1A)".

Observations show that there is a strong, near-linear relationship between spike dry weight at anthesis, $SDW_a$ (g d.wt m-2) and $GN$ (grains m-2). There is secondary variation in this relationship as determined by reproductive efficiency of the spikes to set grains per unit of spike growth, termed the fruiting efficiency, $FE$ (grains per g d.wt SDW). Investigations have demonstrated that, when focusing solely on genotypic influences on $GN$, the $FE$ plays a more important role than $SDW_a$ Slafer et al. (2022). However, there is a stronger relationship between $GN$ and $SDW_a$ under environmental variations (Pretini et al., 2021). There is also some evidence of interactions between genetic and environmental effects, and a degree of compensation between $SDW_a$ and $FE$ as they generally show a negative relationship to one another (Terrile et al., 2017).

##### 7.1.1 Target Grain Production

The grain production module for wheat in DAESIM2 attempts to capture the key factors that control $GY$. The potential grain density, $S_d^{pot}$ (often denoted as grain number, $GN$, grains m$^{-2}$), is calculated using a relatively simple relationship between $FE$ and $SDW_a$, which includes a sigmoid factor to modulate $GN^{pot}$ at the low and high extremes of $SDW_a$, as follows:

$$S_d^{pot} = FE \times SDW_a \times \frac{1}{1 + e^{-k(SDW_a - SDW_{50})}} \tag{57}$$

where $SDW_a$ is the spike dry weight at the start of anthesis (g d.wt m$^{-2}$), $FE$ is the reproductive (fruiting) efficiency of the spikes to set grains per unit of spike growth $FE$ (grains g d.wt$^{-1}$), while $k$ and $SDW_{50}$ determine the sigmoid relationship between $SDW_a$ and $S_d^{pot}$. This formulation ensures that the potential grain number is determined by assimilate supply during the spike formation phase.

Then, the potential grain dry weight, $W_{seed}^{pot}$ (g d.wt m$^{-2}$), is determined by:

$$W_{seed}^{pot} = S_d^{pot} \times GW \tag{58}$$

Where $GW$ is the mean individual weight of the grains (g d.wt grain$^{-1}$). In wheat, variation in $GW$ tends to have a minor effect on yield (e.g. Peltonen-Sainio et al., 2007; Slafer et al., 2014), thus we assume this parameter is fixed for a given genotype.

It is important to note that in DAESIM2-Plant, the spike is not currently treated as a separate organ from the stem. Instead, the development of the stem and spike are modeled in distinct growth phases, with different growing-degree day requirements, allocation coefficients and turnover rates. During the vegetative phase, there is no spike formation so the stem pool corresponds to the stem organ. The subsequent spike formation phase defines a period during which there is enhanced growth, almost entirely contributing to spike formation Fischer and Stockman (see 1980). Thus, in the model, it is assumed that all of the increase in the stem biomass pool corresponds to spike growth, thus spike dry weight ($SDW_a$) in DAESIM2 is inferred as the change in stem biomass during the spike formation phase.

Mathematically, $SDW_a$ is equal to the change in stem biomass from the start of the spike formation and the end of the spike formation (i.e. start of anthesis):

$$SDW_a = W_{stem}(t_0^{spike}) - W_{stem}(t_0^{anthesis}) \tag{59}$$

Where $W_{stem}$ is the stem dry biomass (g d.wt $m^{-2}$), $t_0^{spike}$ is the beginning of the spike formation phase and $t_0^{anthesis}$ is the beginning of the anthesis phase, which (again) assumes that all of the stem biomass gain during the spike formation period is due to spike growth. Environmental effects on plant processes such as photosynthesis and developmental rate, which ultimately determine the stem biomass change during this critical period can therefore $GN^{pot}$ via $SDW_a$.

Note that observed $SDW_a$ typically ranges from 100-250 g d.wt $m^{-2}$, observed seed densities (grain numbers) in wheat range from 2630-17510 grains $m^{-2}$ for spring wheat (2160-18330 when including quartiles) and 2290-19340 grains $m^{-2}$ for winter wheat (2060-19340 when including quartiles), while observed single grain weights range from 31.3-41.6 mg for spring wheat (28.4-43.7 mg when including quartiles) and 37.8-46.9 mg for winter wheat (34.7-47.9 mg when including quartiles) (Peltonen-Sainio et al., 2007). Fruiting efficiency, $FE$, typically ranges from 80-210 grains g d.wt$^{-1}$ (Fig. 7 Terrile et al., 2017).

###### 7.1.2 Grain Filling

The grain filling phase determines whether the potential grain number and dry weight is achieved. During the grain filling phase, the grain is supplied with assimilates from two sources: (i) directly from photosynthesis and (ii) remobilisation of stem reserves of non-structural carbohydrates. Both sources of carbon for grain filling continue at their respective rates until the grain filling phase has concluded or the maximum potential dry weight is achieved. New assimilates are supplied to grain via a fixed allocation coefficient multiplied by NPP. The remobilisation of stem reserves is described below.

Remobilisation of stem reserves for grain filling is modelled using a Michaelis-Menten type relationship assuming that this process is limited by the phloem loading rate, following previous research in trees (Schepper and Steppe, 2010; Trugman et al., 2018). The phloem loading rate ($L$; g C d$^{-1}$) is modelled as follows:

$$L = V_{max,L} \frac{C_{organ}}{K_{M,L} + C_{organ}} \tag{60}$$

with $C_{organ}$ the concentration of remobilizable carbon in the stem (g C g d.wt$^{-1}$), $V_{max,L}$ (g C d$^{-1}$) defining the maximum potential loading rate, and $K_{M,L}$ (g C g d.wt$^{-1}$) the concentration at which the loading rate is 50% of $V_{max,L}$ i.e. the kinetic parameters of the Michaelis-Menten function. To avoid the need for modeling complex source-sink dynamics and tracking the state of non-structural carbohydrates in each plant organ, we adapt this for application in the DAESIM2 model. We assume that the ratio of actual Cstem:Cplant provides a proxy for the remobilizable carbon in the stem. This means that when the actual Cstem:Cplant is high (e.g. in wheat this is usually during spike formation), there is ample remobilizable carbon. Conversely, when the actual Cstem:Cleaf is low, there is less remobilizable carbon in the stem and relatively more structural carbon. Thus, we first calculate the ratio of stem to leaf carbon:

$$C_{organ} \propto R_{S,L} = \frac{C_{stem}}{C_{leaf}} \tag{61}$$

Thus $C_{organ}$ in Eq. 60 is replaced by the ratio Cstem:Cleaf and $K_{M,L}$ now represents the Cstem:Cleaf ratio at which the loading rate is 50% of $V_{max,L}$.

###### 7.1.3 Model Sensitivity Tests

Figure 1 shows how the parameters of the grain production model control the relationship between $SDW_a$, potential $GN$ and potential grain yield, $GY$. These show that the model produces a near-linear relationship between $SDW_a$ and $GN$ when there is adequate levels of $SDW_a$ as has been shown by observations (Fischer et al., 2024; Pretini et al., 2021), while it is non-linear as $SDW_a$ nears zero, to capture the potential for grain production from new assimilates during grain filling, even if spike mass is low at anthesis. It also reproduces the observed linear relationship between FE and GN (see Fig. 7 Terrile et al., 2017) as shown below.

The parameters $k$, $SDW_{50}$ and $GN_{max}$ should be fairly conservative and thus kept relatively constant, with acceptable ranges of $k$=[0.01-0.03], $SDW_{50}$=[80-150] and $GN_{max}$=[18000-20000]. The $FE$ typically ranges between 80-210 grains g d.wt-1, which should be specified per genotype. This may be modified

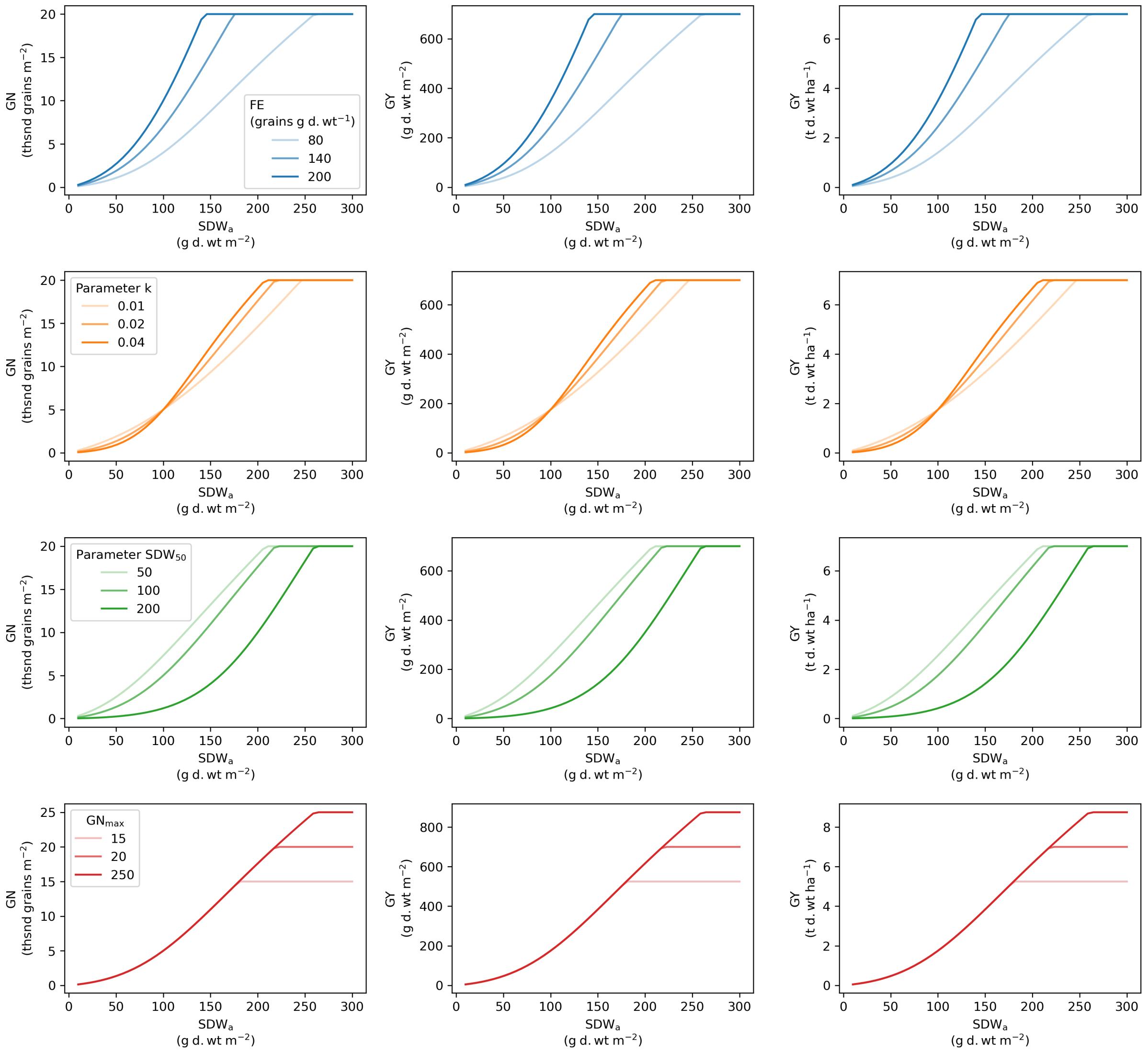


Figure 1: DAESIM2-Wheat grain production model parameter sensitivity tests. Grain yields (GY) in the second and third columns are calculated by multiplying $GN$ by a grain weight of 35 mg.

by environmental conditions in future e.g. excessive heat or frost during grain filling are likely to reduce FE.

#### 7.2 Coupling to Carbon Allocation

Given the potential grain density, $S_d^{pot}$, and thousand kernel weight, $W_{seed}^{TKW}$, we can determine the target (potential) seed biomass pool, $W_{seed}^{pot}$ (g d.wt m-2), as:

$$W_{seed}^{pot} = W_{seed}^{TKW} \times S_d^{pot} \tag{62}$$

which is simply the multiplication of the kernel weight by the grain density ($S_d^{pot}$).

The actual grain biomass pool, $W_{seed}$, will be determined by the amount of biomass flowing into the seed pool over a given time interval i.e. grain filling. This flux should continue until the maximum potential seed pool size, $W_{seed}^{pot}$, is reached. The allocation coefficient from assimilates to the grain pool, $u_G$, is determined as follows:

$$u_G(t) = \begin{cases} u_G^{max}, & \text{if } W_{seed}(t)/W_{seed}^{pot} \leq 1 \\ 0, & \text{otherwise} \end{cases} \tag{63}$$

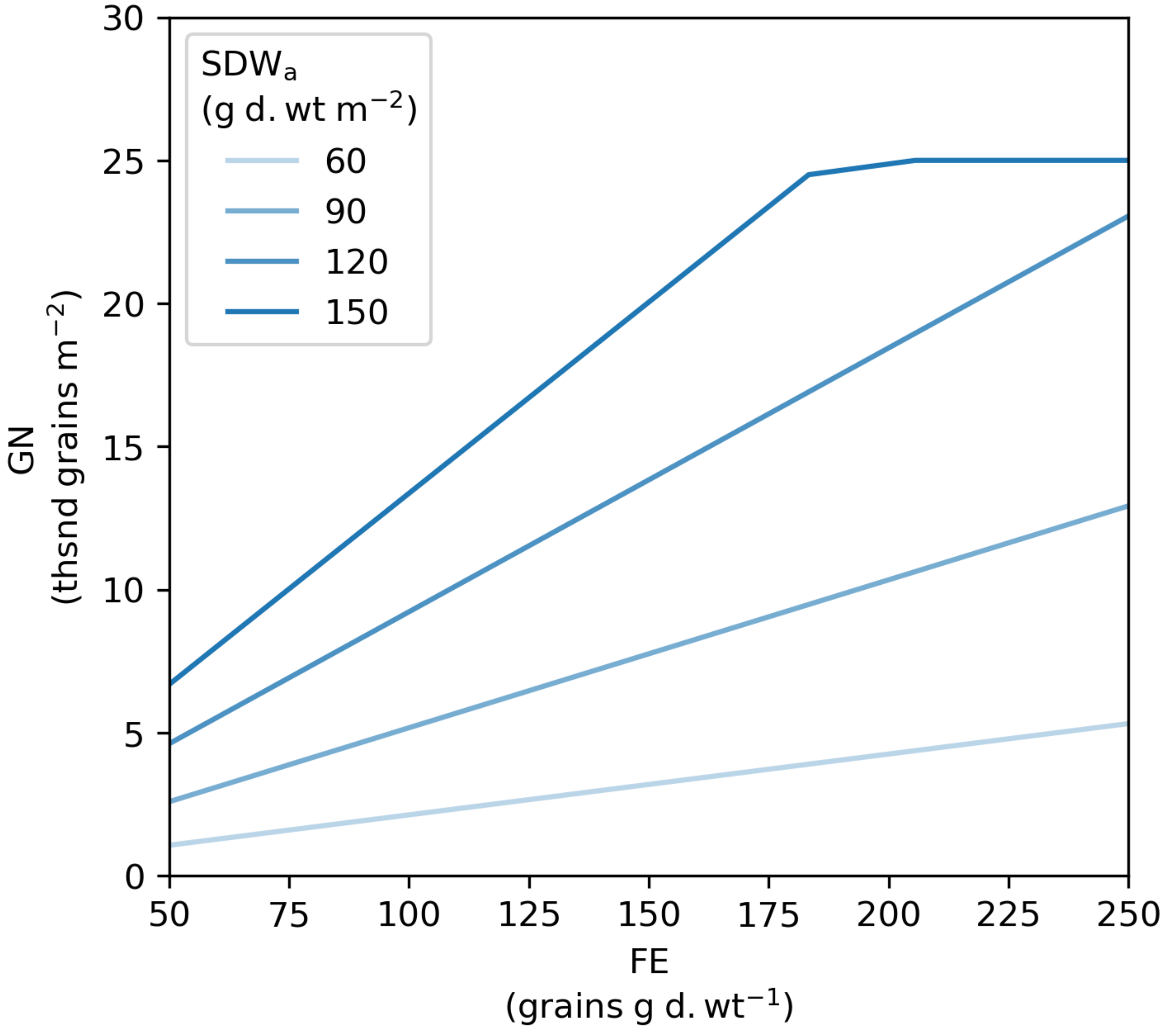


Figure 2: Simulations of DAESIM2 relationships between fruiting efficiency, $FE$, and grain number, $GN$. The lines show isolines for $SDW_a$ at values of 60, 90, 120 and 150 g d.wt m$^{-2}$. The DAESIM2 model simulation uses the following parameter values: $k$=0.03, $GN_{max}$=25, $SDW_{50}$=80. These relationships are consistent with observations in Terrile et al. (2017).

As long as the current seed biomass pool is below the potential, then the flux of carbon continues at the maximum potential rate. The carbon allocation flux is determined as:

$$F_C^{seed}(t) = NPP(t) \times u_G(t) \tag{64}$$

or, if we include stem NSC remobilisation:

$$F_C^{seed}(t) = NPP(t) \times u_G(t) + L(t) \tag{65}$$

Where $L$ is the NSC phloem loading rate from the stem to the seed pool.

This formulation accounts for the effects of assimilate supply on grain filling, as the actual allocation flux to the seed pool during grain filling is limited by source availability, as observed in multiple studies (Kirkegaard et al., 2018; Zhang and Flottmann, 2018).

DISCUSSION: Here we present the second version of our Dynamic Agro-Ecosystem Simulator plant module (DAESim2-Plant). DAESim was designed as a deep process model to be driven with a range of key environmental inputs. It has both biophysical water and carbon flux through leaves and roots and dynamic allocation to stems and grains. DAESim2 also has development where the model accumulates daily environmental units until internal thresholds are met and phase changes occur including the critical transition to flowering that shifts carbon to stems and seeds. We designed DAESim to be constrained by known physiological limits such as light and water, rather than needing calibration across a large range of environments. In this way DAESim could estimate biomass growth and yield as well as below ground allocation, for remotely sensed paddocks in conditions far from regional trial sites.

Another goal is to model generic plant functional types, with subtle changes for different species or genotypes, rather than attempt to calibrate a different model for each of dozens of common cultivars each for dozens of species. As shown, DAESim2 has different stages and allocations for wheat and canola. We envision parameter alterations to adjust for key traits like winter vs spring flowering or short vs adaptive root lengths when trials contrasting these morpho-types are available. However, multi parameter combinations to fit complex yield and biomass allocation differences among large breeding populations grown under homogeneous conditions was not our goal. We rely on the compounding physiological process dependencies to generate the heterogeneous outputs when driven by modeled spatial and temporal field environments seen within and among agricultural growing regions.

In future, new functional types such as perennial grasses, C4, legumes, and woody trees can be incorporated into DAESim2. Together this suite of functional (geno)types can respond to environment (light, temperature, soil moisture) and management (planting time/density, harvest time/intensity) such that their separate and join effects and be partitioned, so called GxExM. With DAESim, it is actually the dynamic daily local environmental inputs triggered by the management intervention and the functional genetic processes that give the ultimate biomass and yield outputs, or ExMxG.

Ultimately our goal is to estimate the full agro-ecosystem water and carbon cycle across mixed functional plant types to include crop rotations, perennial pastures and tree rows responding to dynamic environment and management including cover crops, rotational grazing and tree thinning. Below ground carbon allocation is explicit and outputs from 1 year can be carried into the next as soil environmental inputs providing interannual accounting and compounding benefits. DAESim aims to be a tool model and monitor adaptive management under changing and challenging conditions and to guide on ground validation measurements when and where is most appropriate. It could give confidence to farmers, financiers and regulators aiming to reward best practices with rigor, transparency and more certainty than current approaches.
